# Polar targeting and assembly of the *Legionella* Dot/Icm type IV secretion system (T4SS) by T6SS-related components

**DOI:** 10.1101/315721

**Authors:** KwangCheol C. Jeong, Jacob Gyore, Lin Teng, Debnath Ghosal, Grant J. Jensen, Joseph P. Vogel

**Affiliations:** Department of Molecular Microbiology, Washington University School of Medicine, St. Louis, MO 63110, USA; Department of Animal Sciences &, Emerging Pathogens Institute, University of Florida, Gainesville, FL 32611, USA; Division of Biology and Biological Engineering, California Institute of Technology, Pasadena, CA 91125, USA; Howard Hughes Medical Institute, California Institute of Technology, Pasadena, CA 91125, USA

## Abstract

*Legionella pneumophila*, the causative agent of Legionnaires’ disease, survives and replicates inside amoebae and macrophages by injecting a large number of protein effectors into the host cells’ cytoplasm via the Dot/Icm type IVB secretion system (T4BSS). Previously, we showed that the Dot/Icm T4BSS is localized to both poles of the bacterium and that polar secretion is necessary for the proper targeting of the *Legionella* containing vacuole (LCV). Here we show that polar targeting of the Dot/Icm core-transmembrane subcomplex (DotC, DotD, DotF, DotG and DotH) is mediated by two Dot/Icm proteins, DotU and IcmF, which are able to localize to the poles of *L. pneumophila* by themselves. Interestingly, DotU and IcmF are homologs of the T6SS components TssL and TssM, which are part of the T6SS membrane complex (MC). We propose that *Legionella* co-opted these T6SS components to a novel function that mediates subcellular localization and assembly of this T4SS. Finally, in depth examination of the biogenesis pathway revealed that polar targeting and assembly of the *Legionella* T4BSS apparatus is mediated by an innovative “outside-inside” mechanism.

## Introduction

*Legionella pneumophila* is a Gram-negative bacterium that is the causative agent of Legionnaires’ disease, a fatal form of pneumonia in elderly and immunocompromised individuals. This bacterium grows within a variety of freshwater protozoa in the environment and is transmitted to humans by inhalation of contaminated aerosols^1^. Human disease is caused by the ability of *L. pneumophila* to survive and replicate inside alveolar macrophages^2–4^. Upon uptake into phagocytic cells, the bacteria avoid the host endocytic pathway and recruit early secretory vesicles to the phagosome, forming a unique intracellular compartment termed the replicative phagosome, which is the site of bacterial replication^5^. Upon conclusion of the replicative cycle, the bacteria lyse the host cell and spread to uninfected cells.

Replication of *L. pneumophila* within host cells is strictly dependent on the Dot/Icm type IVB secretion system (T4BSS)^6,7^. The Dot/Icm apparatus is made up of 27 components and injects approximately 300 protein effectors into host cells during infection^8,9^. The apparatus comprises multiple subcomplexes. A major subcomplex, termed *Legionella* core-transmembrane subcomplex (LCTM), consists of at least five proteins (DotC, DotD, DotF, DotG, DotH) and spans the periplasm from the inner membrane to the outer membrane^10^. DotC and DotD, two outer membrane lipoproteins, are known to mediate the outer membrane association of DotH similar to pilotins and secretins^11^. DotF and DotG, two inner membrane proteins, link components of the inner membrane to the outer membrane complex of DotC, DotD and DotH^10^. In addition to the LCTM subcomplex, the Dot/Icm apparatus has at least one additional subcomplex that spans the inner membrane called the *Legionella* type IV coupling protein (LT4CP) subcomplex^12–15^.

Over the years, several laboratories have observed a number of Dot/Icm secreted substrates retained on the cytoplasmic face of the phagosomal membrane, adjacent to the poles of the bacteria^16–21^. Consistent with this observation, many Dot/Icm components have been shown by immunofluorescence to be at both poles^21,22^. Moreover, intact apparatus were recently detected at the *L. pneumophila* cell poles by electron cryotomography (ECT)^23^. We then went on to show that polar localization of this T4SS is critically important for this pathogen as polar secretion of Dot/Icm substrates is required for *L. pneumophila* to alter the host’s endocytic pathway, an essential trait for intracellular replication^21^.

In this manuscript, we focus our attention on the biogenesis of the *L. pneumophila* T4BSS, specifically how the Dot/Icm apparatus is targeted to the bacterial poles. We report that two Dot/Icm proteins, DotU (also known as IcmH) and IcmF, are required for the polar localization of the Dot/Icm apparatus. DotU and IcmF are both necessary and sufficient for polar targeting as they are able to target to the poles on their own and are capable of mediating localization of the core-transmembrane subcomplex to the poles in the absence of other Dot/Icm components. In addition to performing a role in targeting, DotU and IcmF are required for the outer membrane association of DotH, one of the first steps in the biogenesis of this fascinating type IV secretion system.

## Results

### The *Legionella* type IV core-transmembrane subcomplex is not capable of targeting to the bacterial poles by itself

Previously we documented that three components of the core-transmembrane subcomplex, DotF, DotG, and DotH, localize to the bacterial poles^21^. Since this subcomplex also contains the lipoproteins DotC and DotD^10^, we examined their localization to confirm they were also present at the poles. As our DotC and DotD antibodies did not function by immunofluorescence, we employed fusions of each protein to the hemagglutinin (HA) epitope tag. Each fusion was functional as it was able to complement the intracellular growth defect of the corresponding deletion strain (Supplementary Fig. 1). Similar to the polar targeting of DotF, DotG, and DotH, we were able to detect DotC-HA and DotD-HA by immunofluorescence at both extremities of early stationary phase grown bacterial cells (Fig. 1A). Demonstrating specificity, no signal could be detected for each antibody in a strain that did not express any of the *dot/icm* genes (we refer to this strain as the “super *dot/icm* deletion” strain or “S∆” strain). Although fluorescence was observed when individual components were expressed in the S∆ strain, the proteins were found at non-polar parts of the cell (Fig. 1A). For each strain, expression of the correct protein(s) was confirmed by western analysis (Supplementary Fig. 2). Based on these observations, we concluded that polar localization of the core subcomplex is likely dependent on the presence of one or more additional Dot/Icm proteins.

**Fig. 1A.**
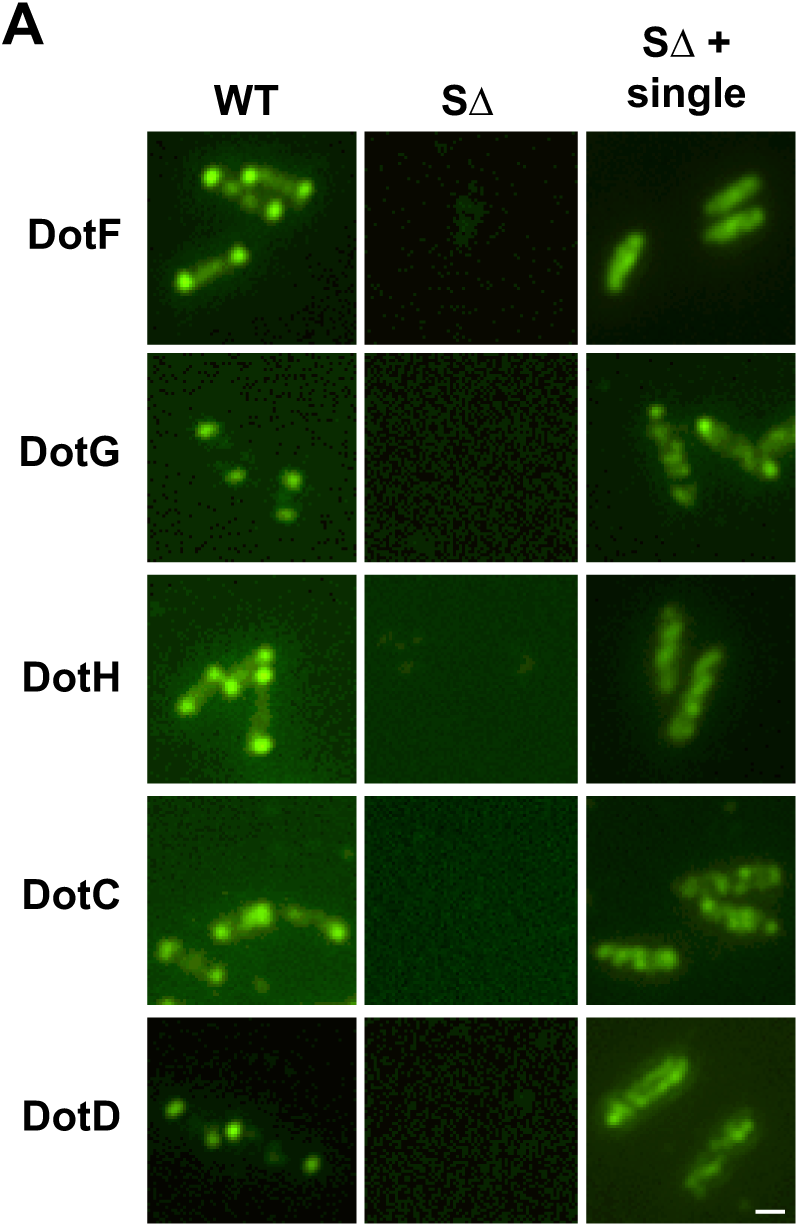
The *Legionella* core-transmembrane subcomplex does not localize to the bacterial poles in a strain lacking the other Dot/Icm proteins (S∆). Broth grown *L. pneumophila* cells were probed with primary antibodies (polyclonal for DotH, DotG, and DotF or HA monoclonal for DotD-HA and DotC-HA), decorated with secondary antibody conjugated with Oregon green and imaged with fluorescence microscopy. Samples assayed include the wild-type strain Lp02 (WT), a strain lacking all 26 *dot/icm* genes (S∆), and the S∆ strain expressing individual core-transmembrane subcomplex components. Imagining of DotC and DotD was done by expressing DotC:HA and DotD:HA in place of the wild-type proteins. Representative images are shown from at least three independent experiments. (Scale bar: 2 μm)

### DotH, DotG, and DotF localization in individual *dot/icm* deletion strains

To determine how the Dot/Icm core-transmembrane subcomplex is targeted to the bacterial poles, we assayed the effect of individual *dot/icm* deletions on the localization of the DotH, DotG, and DotF proteins. As previously reported^21^, DotH staining was observed primarily at both bacterial poles in the majority of wild type (WT) cells (greater than 95% in 300 scored cells) (Fig. 1B and Supplementary Fig. 3). The staining was specific to DotH, since no fluorescence staining was observed in the Δ*dotH* mutant (yellow boxed in Fig. 1B). Interestingly, the majority of the *dot/icm* mutants did not appear to significantly affect DotH localization. However, the staining pattern for DotH was noticeably altered in four mutants including the Δ*dotC*, Δ*dotD*, Δ*dotU* and Δ*icmF* mutants (red boxed in Fig. 1B).

**Fig. 1B.**
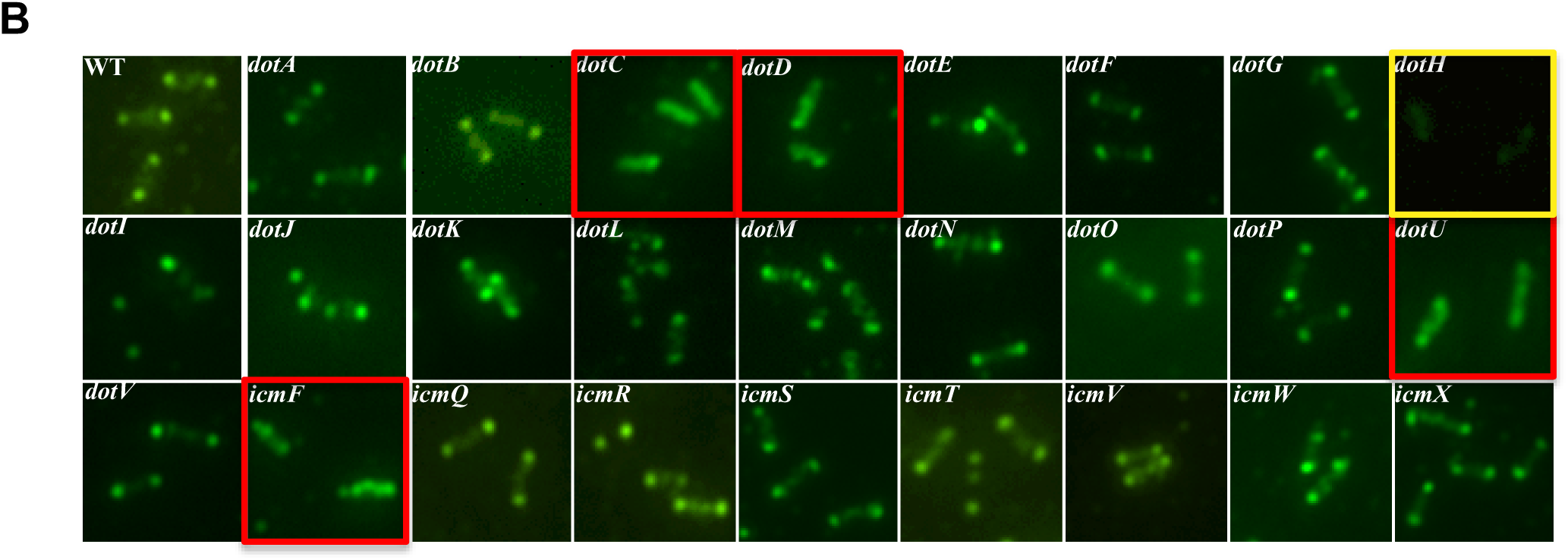
DotH localization in wild-type *Legionella* and in individual *dot/icm* deletions. DotH was detected by immunofluorescence microscopy in wild-type cells (WT) and individual *dot/icm* mutant strains. The corresponding deletion strains is boxed in yellow and *dot/icm* deletions that have an effect are boxed in red. Representative images are shown from at least three independent experiments.

Having seen an effect on DotH localization, we then examined targeting of two other components of the LCTM subcomplex, DotG and DotF. A DotG and a DotF-specific signal could be detected in WT cells but was missing in the corresponding deletion strains (yellow boxed in Supplementary Fig. 4). Interestingly, the same four mutants described above (Δ*dotC*, Δ*dotD*, Δ*dotU* and Δ*icmF*) also affected the polar localization of DotG and DotF (red boxed in Supplementary Fig. 4). The one notable difference for DotF was that its targeting was also dependent on the outer membrane protein DotH. As a result, these data reveal both an intra-dependency between components of the LCTM subcomplex for polar localization and a role for DotU/IcmF in the proper localization of the LCTM subcomplex.

### Mislocalization of DotH, DotG, DotH to non-polar punctae

Although DotH, DotG, and DotF do not target to the poles in the absence of *dotC*, *dotD*, *dotU* or *icmF*, we were surprised that they appeared to localize in a pattern reminiscent of cytoplasmic staining. This was unexpected as we predicted the proteins, upon de-localization from the poles, would remain associated with the cell envelope and display a ring-like pattern consistent with their presence in the inner membrane, periplasm or outer membrane^21^. However, it was difficult to conclusively assign a location to the proteins because of an unexpected marked increase in the fluorescent signal of each protein when compared to that observed in wild-type cells (Fig. 1B). The enhanced immunofluorescence signal was not caused by an increased amount of protein as shown by Western analysis (Supplementary Fig. 5) but could possibly be due to enhanced exposure of epitopes in certain mutant backgrounds, thus obscuring their actual location within the cells.

In order to decipher their true location, the position of the three proteins was re-examined using decreased amounts of the primary antibodies and deconvolution microscopy. Under these conditions, it became readily apparent that DotH, DotG, and DotF proteins were not cytoplasmically localized in a ∆*dotU*∆*icmF* (∆UF) mutant or a ∆*dotC* mutant but instead appeared to consist of specific, mostly non-polar, punctae (Fig. 2A and Supplementary Fig. 6). Based on the location of the puncta, we initially considered they might be in a spiral or a helix similar to what was originally reported for YFP-MreB, even though this was shown to be artificially generated due to the YFP fusion^24^. In contrast, the altered localization of DotH, DotG, and DotF was not artificially generated as we localized non-fused, wild-type Dot proteins expressed from their native positions on the chromosome. The Dot punctae in the mutants did not co-localize with the YFP-MreB helices (Fig. 2B) and careful examination revealed they were not actually spirals/helices as these patterns would wrap around a cell with one diagonal pitch on the “top” of the cell and the opposite pitch on the “back” side, which did not occur. Nevertheless, it is clear that the Dot punctae are not primarily at the poles and may co-localize with a specific, unknown subcellular structure.

**Fig. 2.**
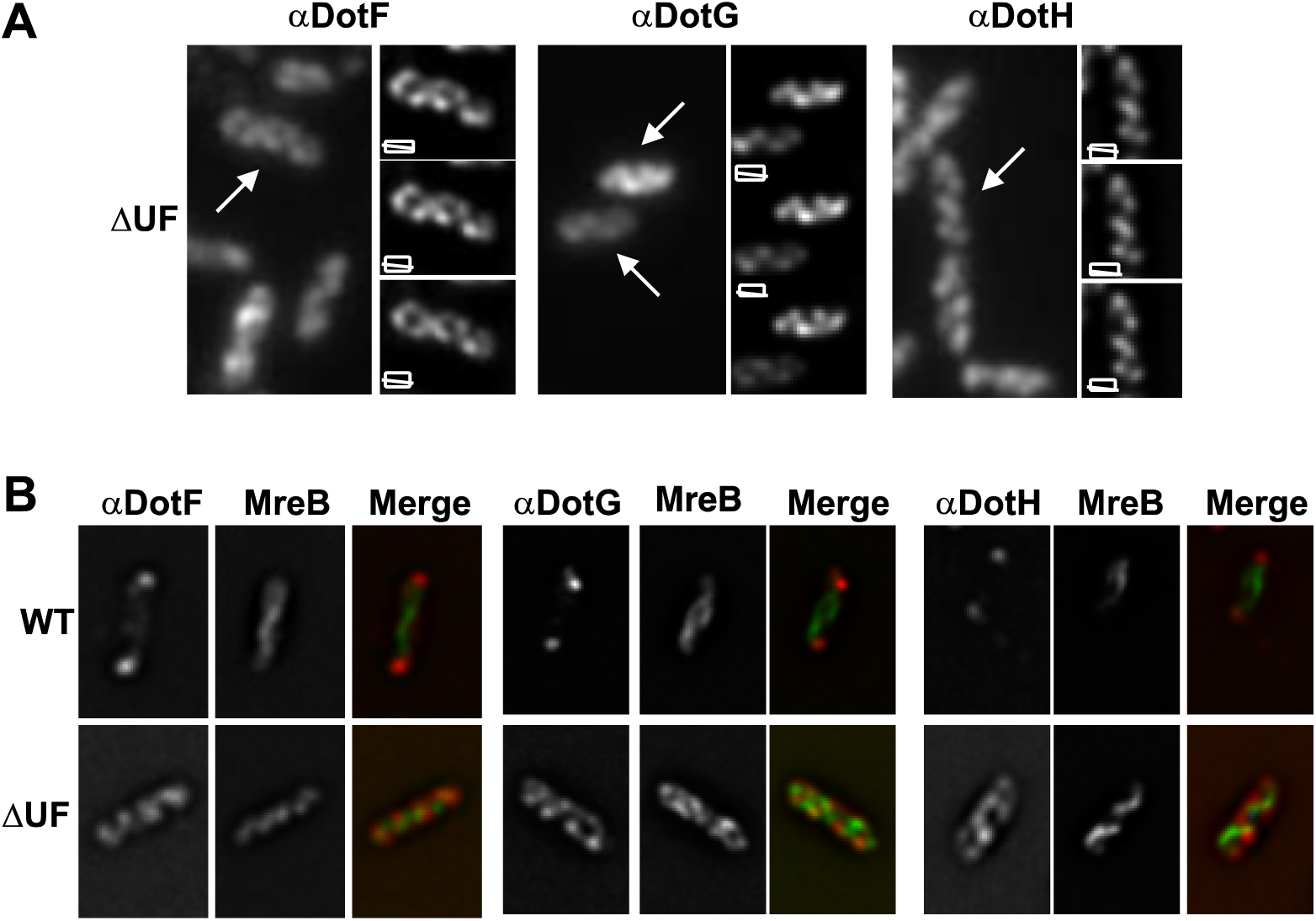
DotH, DotF and DotG proteins are found in non-polar punctae in the ∆*dotU* ∆*icmF* mutant strain. (A) DotH, DotG, and DotF localization was assayed using a lower amount of primary antibody and deconvolution microscopy. For each strain, four images are shown including an optical section before deconvolution (left) and three images of a top optical section, a middle optical section, and a bottom optical section (right panel). Arrows in the left image indicate the cell image that was deconvoluted. (B) Localization of DotH, DotG, and DotF in wild type cells (WT) and ∆*dotU* ∆*icmF* mutant strain expressing YFP-MreB. For each protein, images are shown for the corresponding Dot antibody, YFP-MreB, and a merged image. Representative images are shown from at least three independent experiments.

### DotU and IcmF serve as anchor proteins for the Dot/Icm T4SS polar localization

Based on these results, we hypothesized that DotU/IcmF may function directly as an anchor or a landmark to recruit the LCTM subcomplex to the bacterial poles. This premise predicts that DotU and IcmF should be able to localize to the poles of a wild-type cell in the absence of other components of the Dot/Icm complex. To test this theory, we examined the localization of DotU and IcmF in wild-type cells and in the super *dot/icm* deletion strain expressing only DotU and IcmF, indicated as “S∆(UF)”. Consistent with our prediction, polar localization of DotU and IcmF was observed in greater than 90% of the wild-type cells (Fig. 3A and Supplementary Fig. 7A). The detected fluorescence was specific to the proteins as no signal was observed in a strain lacking *dotU* and *icmF* or in the S∆ strain (Fig. 3A). Most notably, expression of only DotU and IcmF in the S∆(UF) strain allowed detection of the proteins specifically at the poles and at levels indistinguishable from the wild-type strain (Fig. 3A). Thus, DotU and IcmF fulfill the criteria as anchors for the recruitment of the LCTM subcomplex to the bacterial poles.

**Fig. 3.**
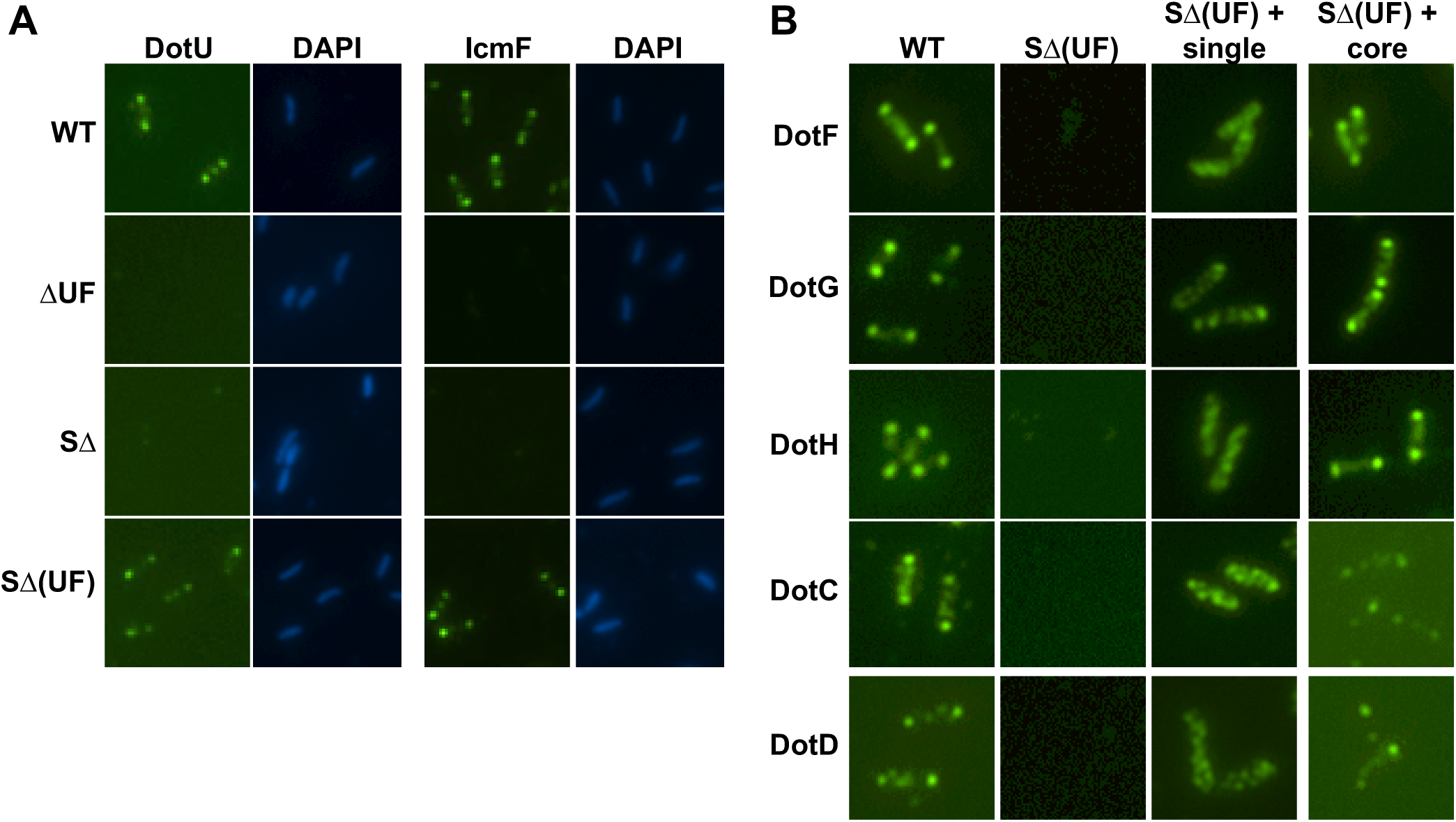
DotU and IcmF localize to the bacterial poles in the absence of the *Legionella* T4SS. (A) DotU and IcmF localization was assayed in the wild-type strain Lp02 (WT), ∆*dotU* ∆*icmF* mutant strain (JV1181), the super *dot/icm* deletion strain (S∆, JV4044) and the S∆ strain expressing *dotU* and *icmF* from the chromosome (SΔ(UF), JV5319). Shown are Dot staining (left) and DNA stained with DAPI (right) for each set. (B) Localization of Dot proteins was assayed in the wild-type strain (WT), the S∆ strain encoding *dotU* and *icmF* (S∆(UF)), the S∆(UF) strain expressing individual components of the core-transmembrane subcomplex (S∆(UF) + single), and the S∆(UF) strain expressing all five core components (S∆(UF) + core). Antibodies used for immunofluorescence are indicated to the left of the panels. Representative images are shown from at least three independent experiments.

To confirm the targeting properties of DotU and IcmF, we then examined the localization of the core-transmembrane subcomplex when its components were expressed in the S∆(UF) strain. DotU and IcmF were insufficient to target the five proteins to the poles at wild-type levels when they were expressed individually in the S∆(UF) strain (Fig. 3B, third column), consistent with our data showing a co-dependence of the subcomplex for polar localization. However, the presence of DotU and IcmF remarkably restored polar localization when all five proteins were expressed in the S∆(UF) strain (Fig. 3B and Supplementary Fig. 7B and 8). In summary, DotU and IcmF are able to target to the poles of a *Legionella* strain that does not express any other Dot/Icm proteins and expression of DotU and IcmF is sufficient to localize the LCTM subcomplex to the ends of the bacterial cell when all five components of the subcomplex are present.

### Biogenesis of the core-transmembrane subcomplex of the Dot/Icm T4SS apparatus

The intra-dependence between components of the LCTM subcomplex for proper polar localization by DotU and IcmF suggested this process is likely to be multifaceted. Therefore, we performed a detailed analysis of polar targeting by expressing combinations of the core-transmembrane subcomplex components in the S∆(UF) strain. Each strain was first assayed by western analysis to ensure that they expressed the correct proteins (Supplementary Fig. 9). As observed in Fig. 3B, DotH was unable to localize to the poles when expressed alone in the S∆(UF) strain (Fig. 4A,B). However, wild-type polar targeting of DotH was restored in eight of the fifteen possible combinations of strains expressing various components of the core subcomplex. Strikingly, the eight positive strains always included DotC and notably DotC alone was sufficient to assist DotU/IcmF in the proper targeting of DotH (Fig. 4A,B). The involvement of DotC was consistent with our prior data indicating that DotH localization was affected by the absence of *dotC* in the wild-type strain (Fig. 1B). Oddly DotD was not required for the localization of DotH in the reconstitution strains (Fig. 4A,B), whereas it was required when assayed in otherwise wild-type cells (Fig. 1B) (see Discussion). The presence of DotF and DotG did not improve polar targeting of DotH in the reconstituted strains (Fig. 4A,B) consistent with the absence of a phenotype on DotH localization in the ∆*dotF* and ∆*dotG* strains (Fig. 1B). Based on this analysis, we can conclude the central player for DotH localization is the lipoprotein DotC as it is both necessary and sufficient for DotU and IcmF to target DotH to the poles.

**Fig. 4.**
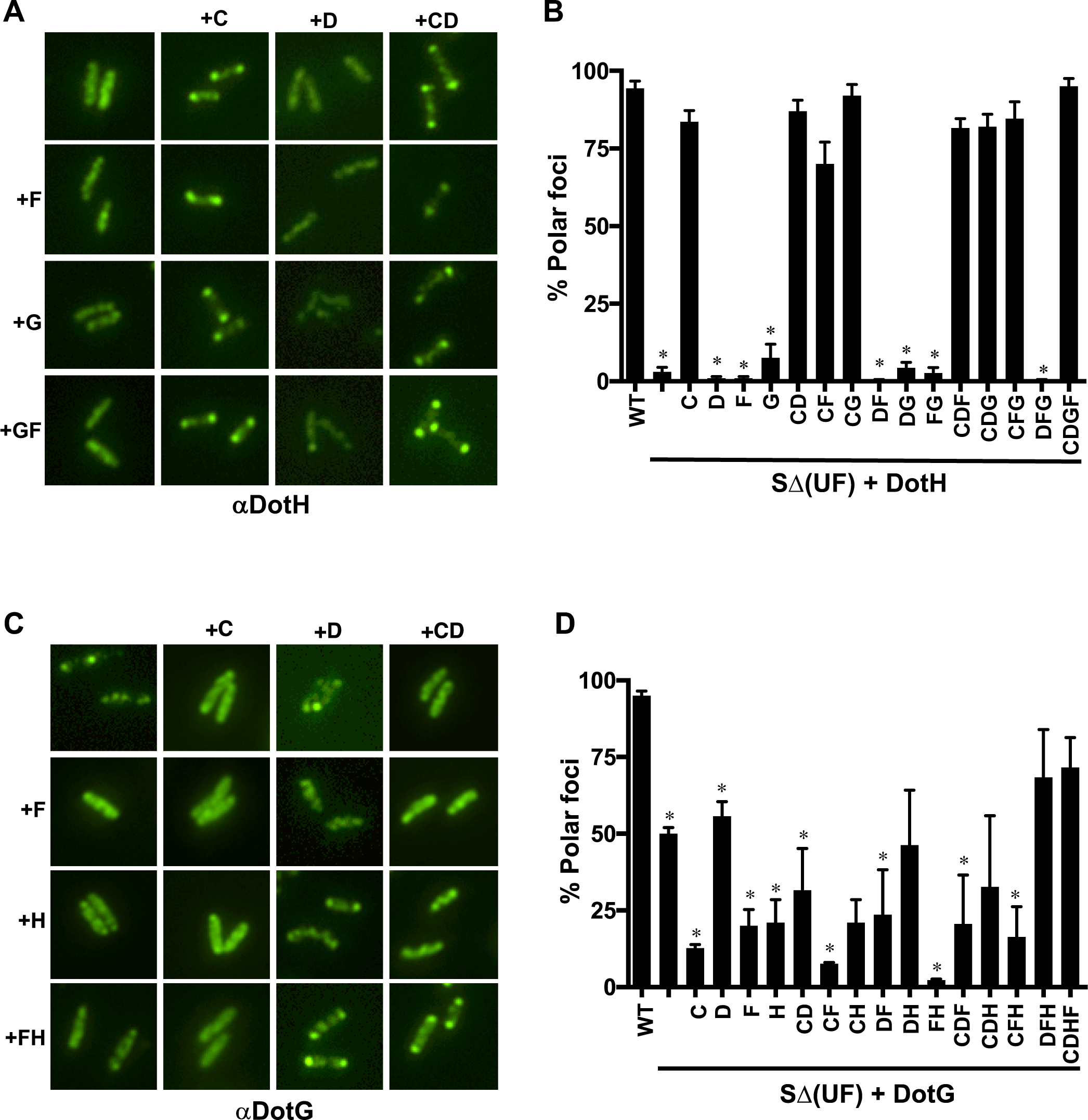
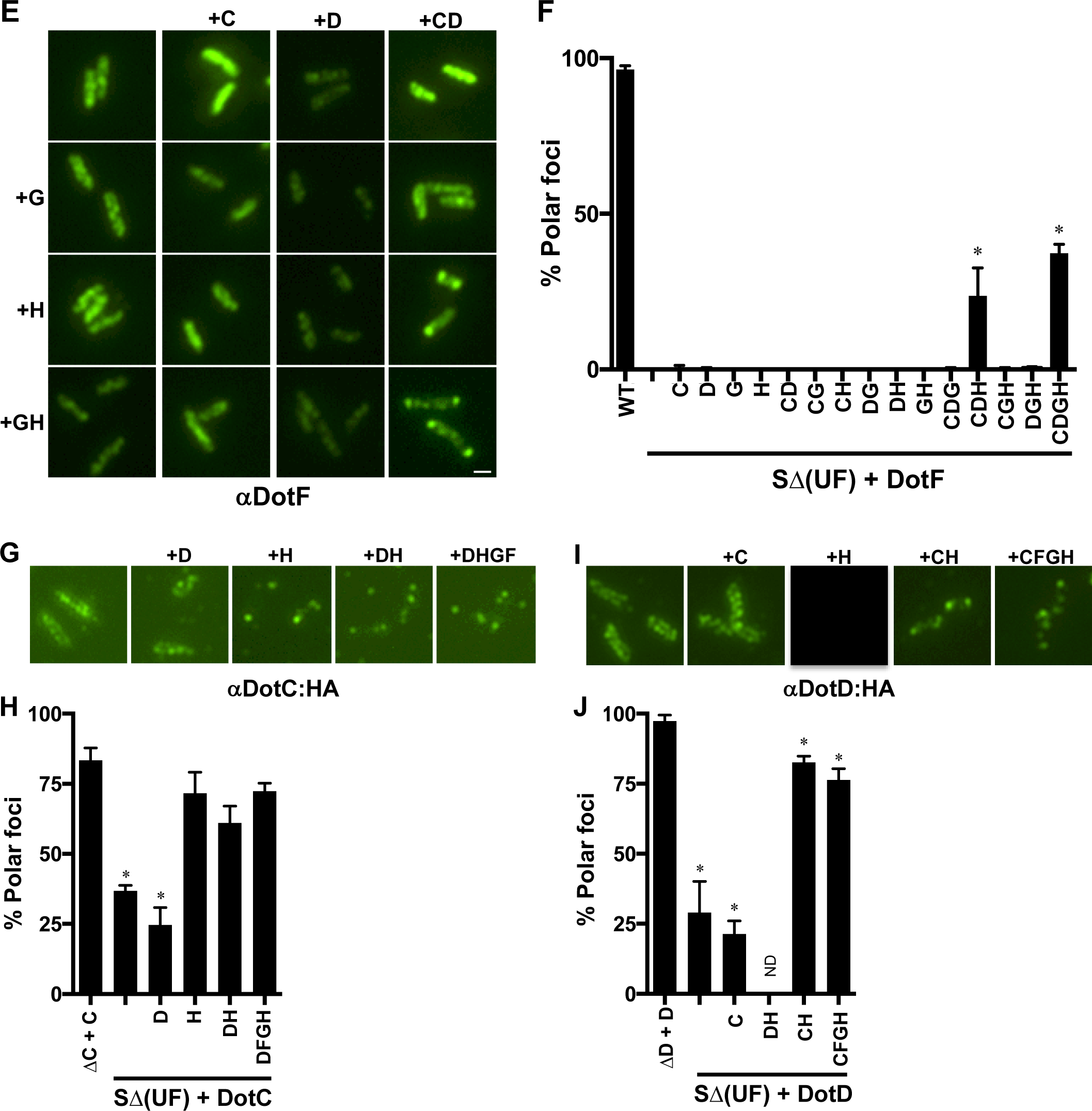
Reconstitution of the core-transmembrane subcomplex in the SΔ(UF) strain. Combinations of the five components of the core-transmembrane subcomplex were expressed in the super *dot/icm* deletion strain encoding *dotU* and *icmF* (S∆(UF)). Representative images for DotH, DotG, DotF, DotC-HA and DotD-HA localization are shown (A, C, E, G, and I, respectively). Proteins expressed in each strain is indicated by labels on the left and top of each panel and the protein localized by IFM is shown below the images. The percent of cells having polar localization of the Dot proteins was determined from three independent experiments (at least 100 cells counted from each experiment) and are shown in panels B, D, F, H and J. Data are presented as means ± SEM. Asterisks indicate statistical difference (* *P*<0.05) compared to the wild-type strain Lp02 (WT) (B, D, and F, respectively) or the corresponding complemented strains (H and J, respectively) by Student’s *t-*test.

In contrast to DotH, DotG localization was somewhat permissive on its own as about half of the cells had protein localized to the poles in the presence of just DotU and IcmF (Fig. 4C,D), although some polar targeting of DotG occurred even in the absence of DotU/IcmF (Fig. 1A). The frequency of DotG being at the bacterial extremities increased in a S∆(UF) strain expressing three other components (DotD, DotF, DotH) or one expressing all five components of the core, approaching the levels observed in the wild-type strain Lp02 (Fig. 4C,D). Enigmatically, DotG localization was sometimes inhibited by the presence of DotC (e.g. compare strains expressing DotG alone vs. DotC & DotG, etc.) (Fig. 4C,D). In terms of DotF, its localization was the most restrictive as partial polar localization was detected in only two of the reconstituted strains (Fig. 4E,F). The presence of DotC, DotD, and DotH restored some polar localization to DotF in the S∆(UF) strain (Fig. 4F). Inclusion of DotG along with the other three components resulted in the best level of DotF localization, although it was significantly less effective than that observed in the wild-type strain Lp02. Thus, localization of DotG was more permissive than DotF but targeting of either protein was optimized in the presence of the other core-transmembrane subcomplex components.

Finally, we examined the location of DotC and DotD in a subset of reconstituted strains, primarily focusing on the lipoproteins and the outer membrane protein DotH. DotC localization did not improve in the presence of the other lipoprotein DotD when compared to a strain expressing just DotC (Fig. 4G,H). In contrast, polar targeting of DotC dramatically increased when co-expressed with just DotH (Fig. 4G,H). As DotH localization reciprocally required DotC (Fig. 4A,B), these results indicate a mutual dependence between the two proteins for polar targeting. In the case of DotD, co-expression of both lipoproteins did not improve the targeting of DotD (Fig. 4I,J) similar to that seen for DotC. We were unable to test if DotH was sufficient for targeting of DotD as DotD protein was not stably found in a reconstituted strain expressing these two proteins (Supplementary Fig. 9). However, the presence of DotC and DotH was sufficient to target DotD to the poles (Fig. 4I,J).

To summarize, we found that DotC and DotH localization occurs first and their targeting is dependent on each other and on the presence of DotU/IcmF. Upon proper localization of DotC and DotH, DotD is recruited to the poles followed by DotG and DotF. Thus, a biogenesis pathway for targeting of the core-transmembrane complex can be inferred to consist of DotU/IcmF going to the poles first, followed by DotC and DotH, then DotD, and finally DotG and DotF (see Discussion).

### Two-step outer membrane association of DotH

Although DotU/IcmF are sufficient to target DotC and DotH to the poles in the S∆(UF) strain, our immunofluorescence assay lacked the resolution to determine if the proteins achieved their final position within the complex. Therefore, we further examined the role of DotU/IcmF in the biogenesis of the Dot/Icm complex by biochemically determining the localization of DotC and DotH via a combination of ultracentrifugation and Triton X-100 solubility. As previously shown^10,13^, the majority of DotH is found in the membrane fraction (M) of the wild-type strain Lp02, although a smaller amount of protein can be detected soluble in the periplasm (S), representing newly synthesized protein prior to its association with a membrane (Fig. 5A and Supplementary Fig. 10). Membrane-associated DotH was not extractable with the detergent Triton X-100, consistent with its linkage with the outer-membrane (O) and not the inner-membrane (I). DotH localization was not dependent on a functional Dot/Icm T4SS complex as it occurred in Lp03, a strain containing a mutation in the inner membrane component DotA (Fig. 5A). Outer membrane association of DotH was dependent on both lipoproteins, DotC and DotD (Fig. 5A). Strikingly, we discovered that DotH outer membrane association did not occur in the ∆*dotU* ∆*icmF* (∆UF) double mutant and this defect could be restored by complementation with a clone expressing *dotU icmF* (UF) (Fig. 5A). Thus, DotH outer membrane linkage requires the lipoproteins DotC/DotD and the targeting factors DotU/IcmF and is consistent with our immunofluorescence data showing that polar localization of the DotH is dependent on the same four proteins.

**Fig. 5AB.**
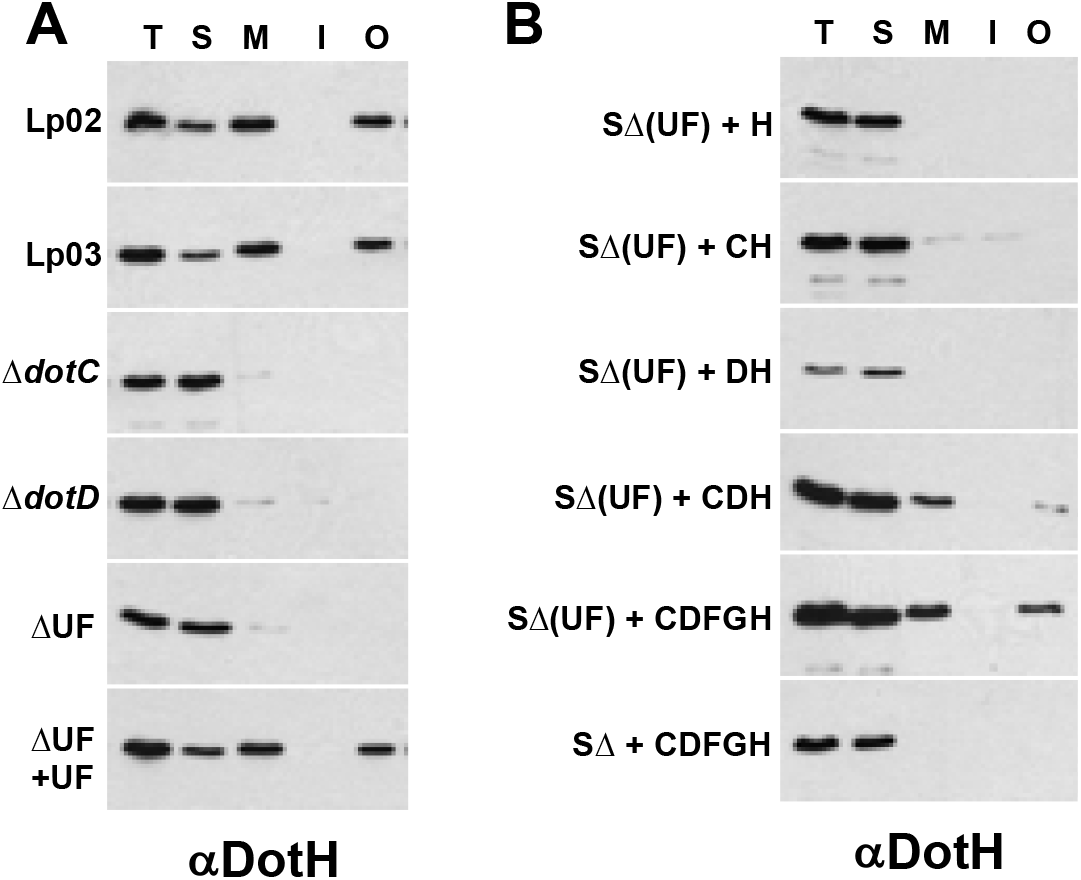
DotH localization to the outer membrane requires the anchor proteins DotU and IcmF. Cells were fractionated by a combination of ultracentrifugation and Triton X-100 solubility, proteins were separated by SDS-PAGE and probed in Westerns using DotH specific antibodies. (A) DotH localization was determined in the wild-type strain Lp02 (WT), the *dotA* mutant Lp03, ∆*dotC* (JV3743), ∆*dotD* (JV3572), Δ*dotU* Δ*icmF* (JV1181), and Δ*dotU* Δ*icmF* + complementing clone (JV1199). (B) DotH localization was determined in the S∆(UF) strain expressing DotH (JV5405), DotC/DotH (JV5458), DotD/DotH (JV5459), DotC/DotD/DotH (JV5460), the core (JV5443) or the core expressed in the S∆ strain without UF (JV5442). Experiments were done in triplicate and representative images are shown.

To further characterize the location and protein interactions of DotH, we repeated the fractionation experiments employing our previously described LCTM subcomplex reconstituted strains (Fig. 4). Expression of DotH alone in the SΔ(UF) strain did not result in outer membrane association of DotH (Fig. 5B). Interestingly, co-expression of DotC in the S∆(UF) strain did not restore outer membrane association of DotH (Fig. 5B), even though it was sufficient for DotC and DotH to co-localize at the bacterial poles (Fig. 4A,B). Proper outer membrane association of DotH in the presence of UF instead required both DotC and DotD and this property improved in a strain expressing all five components of the LCTM (DotCDFGH). Similar to the wild-type strain, DotH did not achieve outer membrane localization in a reconstituted strain lacking DotU and IcmF (Fig. 5B, bottom panel). These results were specific to UF, DotC and DotD as other combinations of proteins failed to properly link DotH to the outer membrane (Supplementary Fig. 10). Thus, DotU/IcmF are able to target DotC and DotH to the poles but DotH is not able to stably associate with the outer membrane in the absence of DotD. This is consistent with recent observations that a periplasmic complex could not be observed by ECT in the absence of DotD but a structure was discernible in a strain expressing DotH, DotC and DotD^25^.

Considering that DotH requires DotU/IcmF to achieve its correct location with the outer membrane, we examined if this could be due to an effect on the lipoproteins DotC and DotD. In these experiments, we localized DotC and DotD using sucrose gradients rather than detergent solubility as all lipoproteins are Triton X-100 soluble unless bound to an outer-membrane, Triton X-100 insoluble protein. In wild-type cells (Lp02), DotC and DotD were found in the same fractions as MOMP, a well characterized *Legionella* outer membrane protein (Supplementary Fig. 11). Their localization did not change in the super deletion strain expressing DotU and IcmF (S∆(UF)) or in the super deletion strain lacking *dotU* and *icmF* (S∆) (Supplementary Fig. 11), indicating the lipoproteins’ outer membrane association was not dependent on DotU/IcmF. Finally, we re-examined the biochemical association(s) of the lipoproteins using Triton X-100. Previously we showed that DotC and DotD were largely resistant to Triton X-100 extraction in the wild-type strain Lp02 but only in the presence of DotH (Fig. 5C,D)^13^. Consistent with this, the majority of both lipoproteins became detergent extractable when expressed by themselves or together in the S∆(UF) strain, indicating they were not bound to an outer membrane associated protein (Fig. 5C,D). However, expression of the two lipoproteins with DotH resulted in a significant fraction of both lipoproteins becoming resistant to the detergent. Markedly, this did not occur in a strain lacking *dotU icmF* even when all five components of the core were expressed (Fig. 5C,D). These results independently confirm both the existence of a complex consisting of DotC, DotD and DotH associating with the outer membrane of *Legionella* and the dependence on DotU and IcmF for its formation.

**Fig. 5CD.**
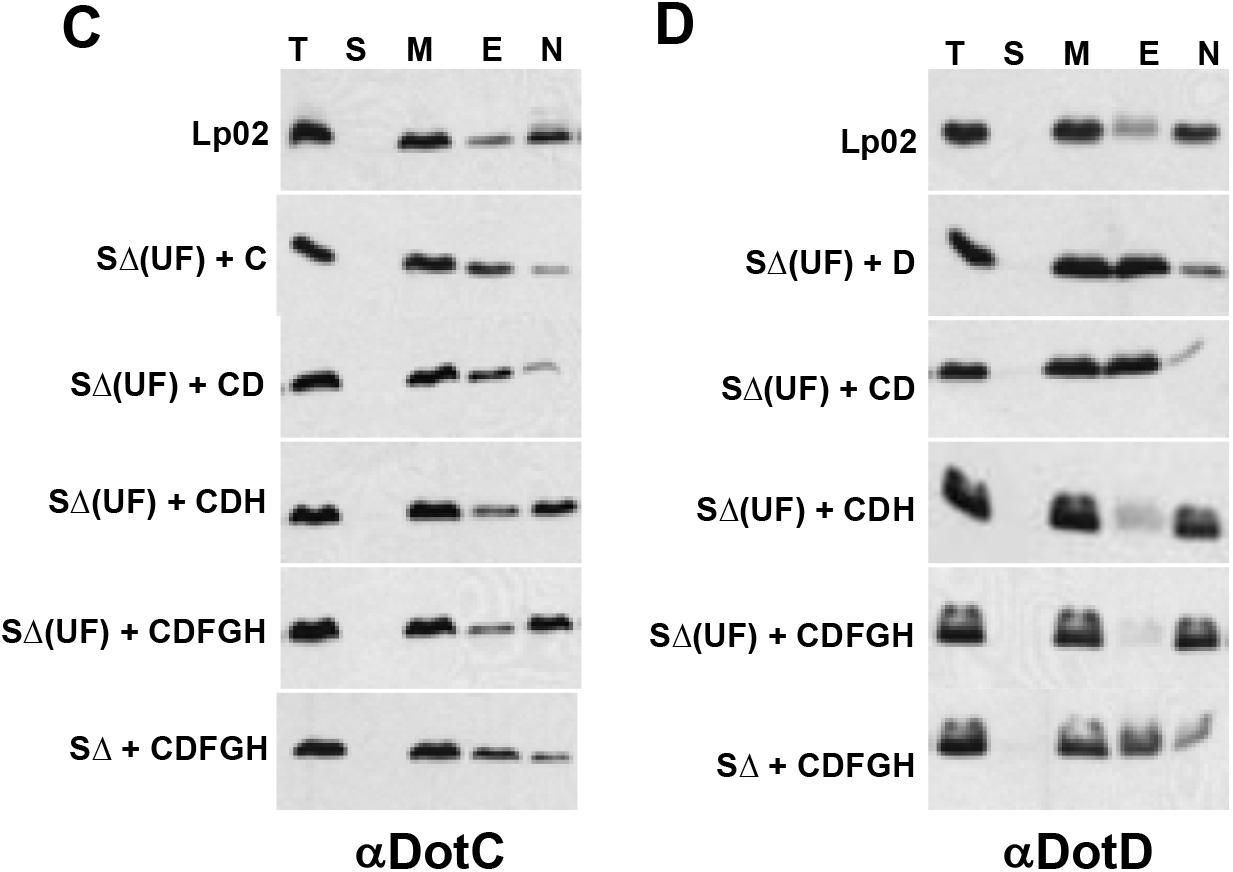
DotC and DotD interaction with DotH in the outer membrane requires DotU and IcmF. Cells were fractionated by a combination of ultracentrifugation and Triton X-100 solubility, proteins were separated by SDS-PAGE and probed in Westerns using DotC and DotD-specific antibodies (A and B, respectively). Interaction of the lipoproteins with DotH was determined in the following strains: wild-type Lp02 (WT), S∆(UF) + DotC (JV5469), S∆(UF) + DotD (JV5470), S∆(UF) + DotC/DotD (JV5471), S∆(UF) strain + DotC/DotD/DotH (JV5460), S∆(UF) + core (JV5463), and the S∆ strain + core (JV5442). Fractions are indicated at the top of the panels and include total proteins (T), soluble proteins (S), membrane proteins (M), Triton X-100 extractable proteins (E), and Triton X-100 non-extractable proteins (N). Experiments were done in triplicate and representative images are shown.

## Discussion

Previously we showed that the Dot/Icm apparatus is at the poles of the bacterial cells because polar secretion must occur in order for this pathogen to alter the endocytic pathway of its host cell. Here we discovered that DotU and IcmF (UF) are necessary and sufficient to serve as targeting factors for the T4SS apparatus. In addition, we found that UF are critical for the initial steps of assembly of the core transmembrane subcomplex, resulting in the stable association of DotH with the outer membrane.

*dotU* and *icmF* encode for inner membrane proteins and are co-dependent for protein stability, indicating they likely function together^26^. ∆*dotU* and ∆*icmF* mutants are severely attenuated for growth in amoebae but are only partially defective for growth in several macrophage lines, unlike most *dot/icm* mutants^26–28^. Based on the observation that several Dot/Icm proteins became destabilized in late stationary phase in the absence of UF, they were hypothesized to function as chaperones and/or assembly factors for the apparatus^26^. In contrast to most Dot/Icm proteins, homologs of DotU and IcmF have not been detected in IncI conjugation systems, the presumed ancestral precursor to the Dot/Icm system^29^. Instead, related proteins are found within T6SSs as DotU and IcmF have homology to T6SS components TssL and TssM, respectively. TssL and TssM are two major factors of the T6SS membrane base plate, which functions as a tail docking station and channel for the passage of the T6SS inner tube^30^. Since no characterized *Legionella* strain possesses an intact T6SS, it is possible that *Legionella* co-opted a version of TssLM when the IncI conjugation system was originally adapted to translocate effectors into host cells. Interestingly, T6SSs are not restricted to the bacterial poles as observed for *Legionella* DotU/IcmF and the Dot/Icm T4SS. So why are DotU/IcmF at the poles in *Legionella*? Perhaps all DotU/IcmF- and TssL/TssM-like proteins target to the poles of *Legionella* due to a unique feature (e.g. lipid componsition) of this bacterial species. Alternatively, DotU and IcmF may have been modified during their co-option by the Dot/Icm system to target to the poles. For example, DotU and IcmF have a different arrangement of transmembrane domains compared to most TssL and TssM proteins and this modification may be responsible for their unique ability to localize to the bacterial poles.

In addition to their role in polar localization, DotU and IcmF also perform a key function(s) in the initial steps of the assembly of the Dot/Icm T4SS. Based on our results, we can now describe a molecular pathway for how Dot/Icm systems begin to assemble. The process begins by the localization of DotU/IcmF to the poles and is followed by the recruitment of both DotC and DotH to the poles. At this stage, the lipoprotein DotC is stably inserted in the outer membrane via its lipid domain whereas DotH remains soluble in the periplasm, presumably loosely associated with DotH and/or IcmF. Next, DotD arrives at the poles and assists in and/or directly mediates the outer membrane association of DotH. After formation of the DotC/DotD/DotH subcomplex, both DotG and DotF are recruited to the poles, thus linking the inner and outer membranes. DotG appears to be able to target to the poles to some level on its own but localizes more efficiently in the presence of other components of the apparatus. DotF localization is strongly dependent on DotC, DotD, and DotH but improves by the presence of DotG. Thus, a biogenesis pathway for targeting of the core-transmembrane complex can be inferred to consist of DotU/IcmF going to the poles first, followed by DotC and DotH, then DotD, and finally by DotG and DotF. A transmission electron microscopy study on the *Legionella* core complex reached similar conclusions for several of the latter steps in the pathway^31^.

This assembly pathway makes sense in light of the recent elucidation of the Dot/Icm T4SS molecular architecture by electron cryotomography (ECT)^25^. In that work, the periplasmic domain of IcmF was found to potentially form part of a “plug” at the heart of the T4BSS midway through the periplasm. The plug is surrounded by a ring of 13 DotC/DotH complexes, which are in contact with a peripheral second ring of DotD proteins, thus generating a defined subcomplex associated with the outer membrane. DotG forms a channel above and below the DotC, DotD, DotH rings and may pass through the DotC/DotH rings. DotF makes up the “wings” surrounding the lower DotG channel and also bind to DotH. This rationalizes why polar localization of DotF is depended strongly on the presence of DotH (Fig. 1B) and why DotF and DotG improved each other’s polar localization, since they interact at the base of the channel.

In addition to these findings, the ECT analysis revealed several additional components (DotK, IcmX, and DotA) that appear to interact with and/or be part of this subcomplex^25^. DotK is a third outer membrane lipoprotein that contributes density on the periphery of the subcomplex near the outer membrane and interacts with DotD, DotH and peptidoglycan, likely stabilizing the complex in the cell envelope^25^. Thus DotK is likely not to be an essential part of the subcomplex and is consistent with the previous observation that *dotK* is dispensable for growth within macrophages^32^ and with the lack of an effect on polar localization of DotH, DotG and DotF in a ∆*dotK* mutant (Fig. 1B). IcmX and DotA form part of the plug at the base of the structure but, similar to the ∆*dotK* mutant, the ∆*icmX* and ∆*dotA* mutants did not have a pronounced effect on polar localization when examined in wild-type cells. However, these three proteins may function to increase the overall efficiency of polar targeting, particularly that of DotF, as it showed the lowest rates in the reconstitution experiments.

In summary, we now understand how many of the periplasmic components of the Dot/Icm apparatus are recruited to the poles and assemble. Nevertheless, many questions remain including how DotU and IcmF were adapted for polar localization, the molecular details of how they recruit DotC and DotH to the poles, whether the assembly of the remaining components/subcomplexes of the Dot/Icm apparatus are dependent on DotU and IcmF and/or whether there are additional independent, targeting factors. Further investigation of how the *Legionella* Dot/Icm T4BSS targets to the poles and assembles will likely provide key information on cellular processes in bacteria and how this pathogen causes disease.

## Acknowledgements

We thank Dr. Ralph Isberg (Tuft’s University) for antibodies that recognize DotF and DotH, Ms. Emily Buford for technical assistance and Dr. Petra Levin (Washington University) for assistance with deconvolution microscopy. We also recognize Dr. Eep Darwin for key suggestions and critical appraisal of this manuscript. This work was funded by NIH grant AI48052 to J.P.V.

## Experimental Procedures

### Strains and cell lines

All bacterial strains are listed in Table 1. *L. pneumophila* strains were cultured in buffered AYE broth or on buffered charcoal yeast extract (CYE) plates. The media were supplemented with 100 μg/ml thymidine as needed (AYET, CYET). All media and antibiotic concentrations for *E. coli* and *L. pneumophila* were used as described previously^18^. The growth phase of liquid cultures was determined by the levels of motility and the optical density.

### Construction of reconstituted plasmids

All used and newly constructed plasmids are listed in Supplementary Table 1. For expression in *Legionella*, all genes were cloned into the vector pJB908 individually or in combination. For new reconstituted plasmids, they were constructed by subcloning of open reading frames of *dotD, dotC, dotF, dotG*, and *dotH*.

### *In vitro* immunofluorescence microscopy

Modified immunofluorescence microscopy (IFM) was carried out as descried previously^33^. In brief, five microliters of an *L. pneumophila* strain, grown to stationary phase culture in broth were briefly fixed in methanol, resuspended in phosphate buffered saline (PBS), and allowed to adhere to poly-L-lysine (Sigma) coated microscope slides. Lysozyme (3 mg/ml in 25 mM Tris-HCl, pH 8.0, 50 mM glucose, 10 mM EDTA) was used to permeabilize the cells, which were then washed with PBS, and incubated with various primary antibodies. After incubation, cells were washed with PBS, decorated with Oregon Green-conjugated goat anti-rabbit IgG, and stained with DAPI to detect DNA. Fluorescence anti-fade reagent was added and the immunostained cells were observed using a fluorescence microscope (Olympus, 100X objective). All images were captured and further analyzed with IPLab software (BD Bioscience).

### Protein fractionations

Protein fractionation was performed by previously described method^10^. In brief, approximately 40 OD600 stationary cells were resuspended in 1 mL of 50 mM Tris-8, 0.5 M sucrose, 5 mM EDTA. After lysozyme treatment, 1.5 mL 50mM Tris pH 8 was added to the samples, and MgSO_4_ was added to 8mM final concentration and sonicated about total 5 min. Samples were centrifuged at 10,000 x g for 15 min at 4° C. Soluble and insoluble proteins were separated by centrifuging 1 mL of this fraction at 100,000 x g for 1 hour. The supernatant from the first spin was re-centrifuged at 100,000 x g for 1 hour to insure complete removal of membrane proteins, and this second supernatant was collected as the soluble protein fraction. Pelleted proteins from the first 100,000 x g spin were resuspended in ice cold 50 mM Tris pH 8 and re-centrifuged at 100,000 x g for 1 hour to remove any contaminating soluble proteins. Total membrane proteins were then resuspended in ice cold 50 mM Tris pH 8. To extract inner membrane proteins, Triton X-100 was added to a final concentration of 1% and the samples were incubated at 37° for 30 minutes. Triton X-100 insoluble proteins were removed by centrifugation of the samples at 100,000 x g for 30 minutes. The supernatants were re-centrifuged at 100,000 x g for an additional 30 minutes to ensure complete removal of Triton X-100 insoluble proteins. Triton X-100 insoluble proteins were resuspended in 1 mL 50 mM Tris pH 8, 1% Triton X-100, and re-centrifuged at 100,000 x g. Triton X-100 insoluble proteins were then resuspended in 1 mL Tris pH 8. Fractions were analyzed by SDS-PAGE followed by immunoblot analysis.

### Statistical analysis

Statistical analysis was performed using Student’s *t*-test of GraphPad Prism 6 (Version 6.0d; GraphPad Software, La Jolla, CA, USA). Data are presented as means ± SEM. Statistical significance was declared if *P*<0.05.

---

**Supplementary Fig. 1.**
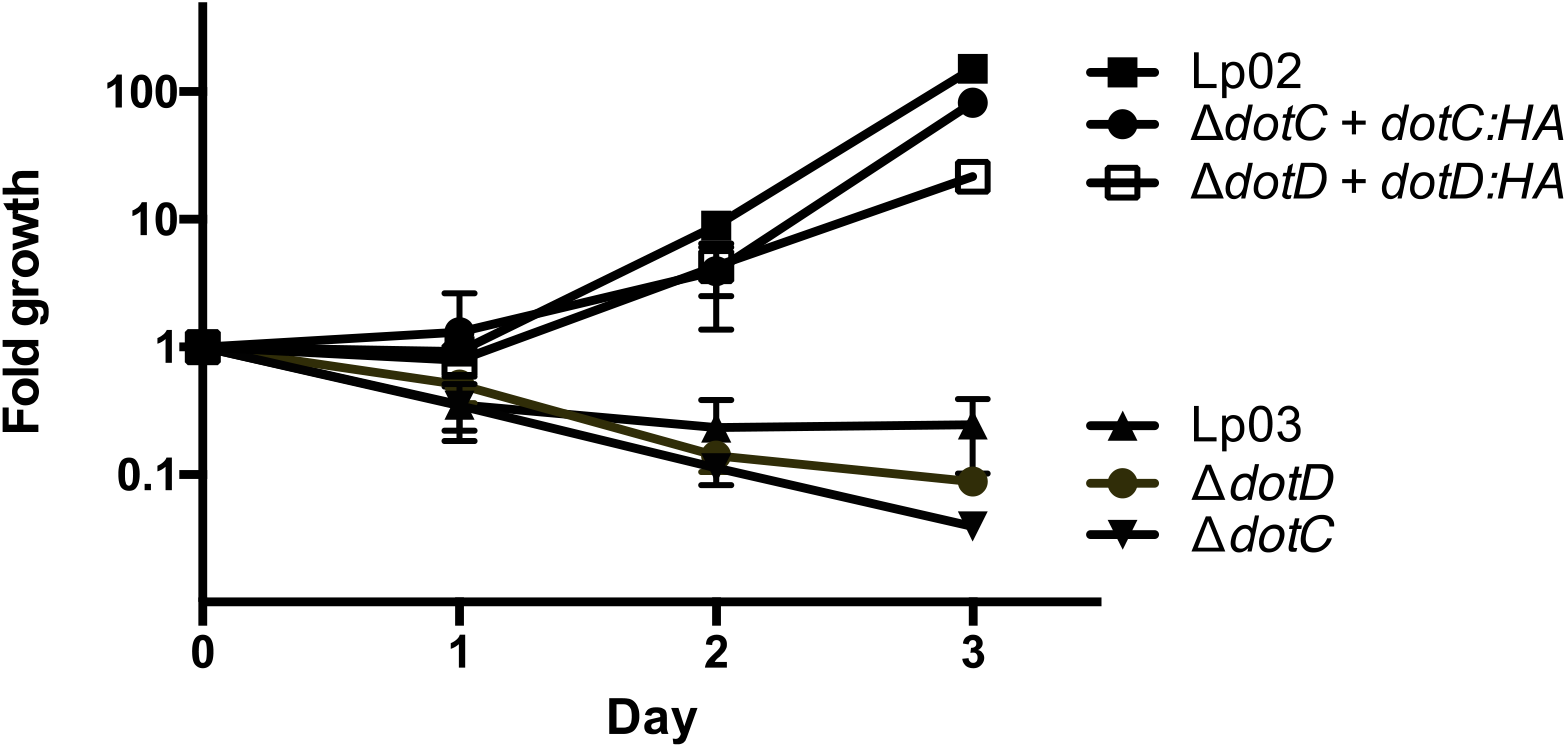
Epitope tagged versions of DotC and DotD are functional for intracellular growth. DotC and DotD were fused to the HA tag at their c-termini and expressed in a ∆*dotC* mutant and a ∆*dotD* mutant, respectively. A wild type *Legionella* strain (Lp02), a *dotA* mutant (Lp03), a ∆*dotC* mutant containing vector (JV5263) or wild type DotC (JV5264) or DotC-HA3X (JV5484) and a ∆*dotD* mutant containing vector (JV5266) or wild type DotD (JV5267) or DotD-HA3X (JV5268) were used to infect U937 cells. Growth was assayed by plating for colony forming units (CFUs) over three days. Data is representative of three independent experiments.

**Supplementary Fig. 2.**
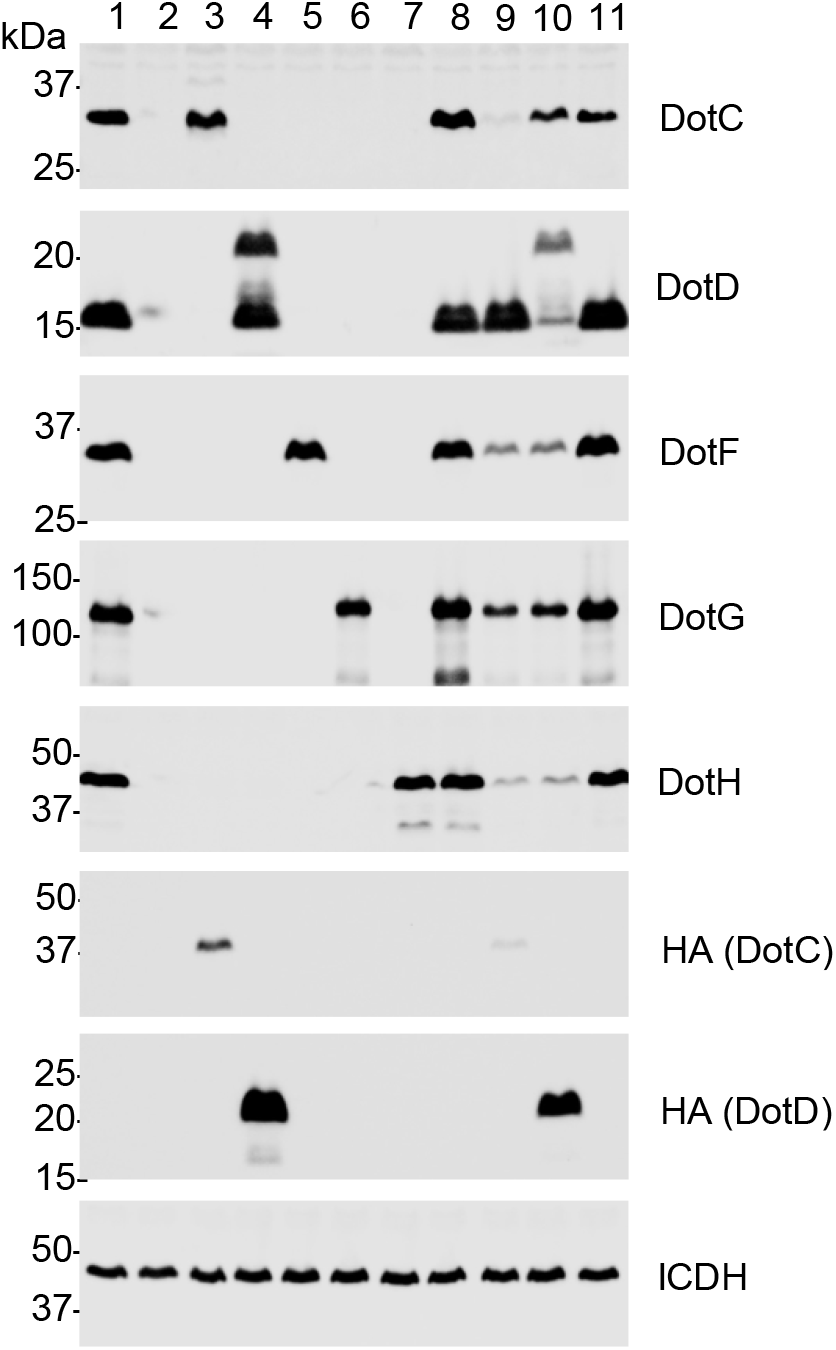
Expression of the correct components in the S∆ strain (JV4044) for Figure 1A. *L. pneumophila* strains were grown to late exponential phase and westerns were done with the indicated antibodies. ICDH, a cytoplasmic housekeeping protein, was used as a loading control. Samples were loaded in the following order: 1. JV1139 (Lp02 + pJB908), 2. JV4209 (S∆ + vector), 3. JV4694 (S∆ + *dotC:HA3x*), 4. JV4695 (S∆ + *dotD:HA3x*), 5. JV4669 (S∆ + *dotF*), 6. JV4688 (S∆ + *dotG*), 7. JV4671 (S∆ + *dotH*), 8. JV5442 (S∆ + *dotCDFGH*), 9. JV9128 (S∆ + *dotC:HA3x dotD dotFGH*), 10. JV9129 (S∆ + *dotD:HA3x dotC dotFGH*), 11. JV1139 (Lp02 + pJB908).

**Supplementary Fig. 3.**
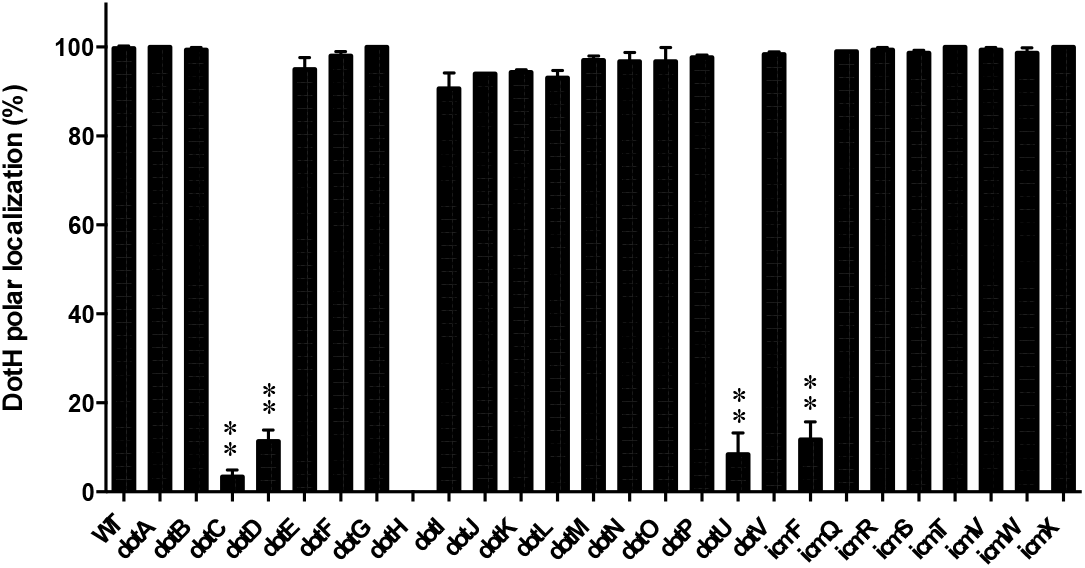
Quantitation of polar localization of DotH for Figure 1B. The percent of cells having polar localization of the DotH in individual *dot/icm* deletions was determined from three independent experiments (at least 100 cells counted from each experiment). Data are presented as means ± SEM. Asterisks indicate statistical difference (** *P*<0.005) compared to the wild-type strain Lp02 (WT) by Student’s *t-*test.

**Supplementary Fig. 4AB.**
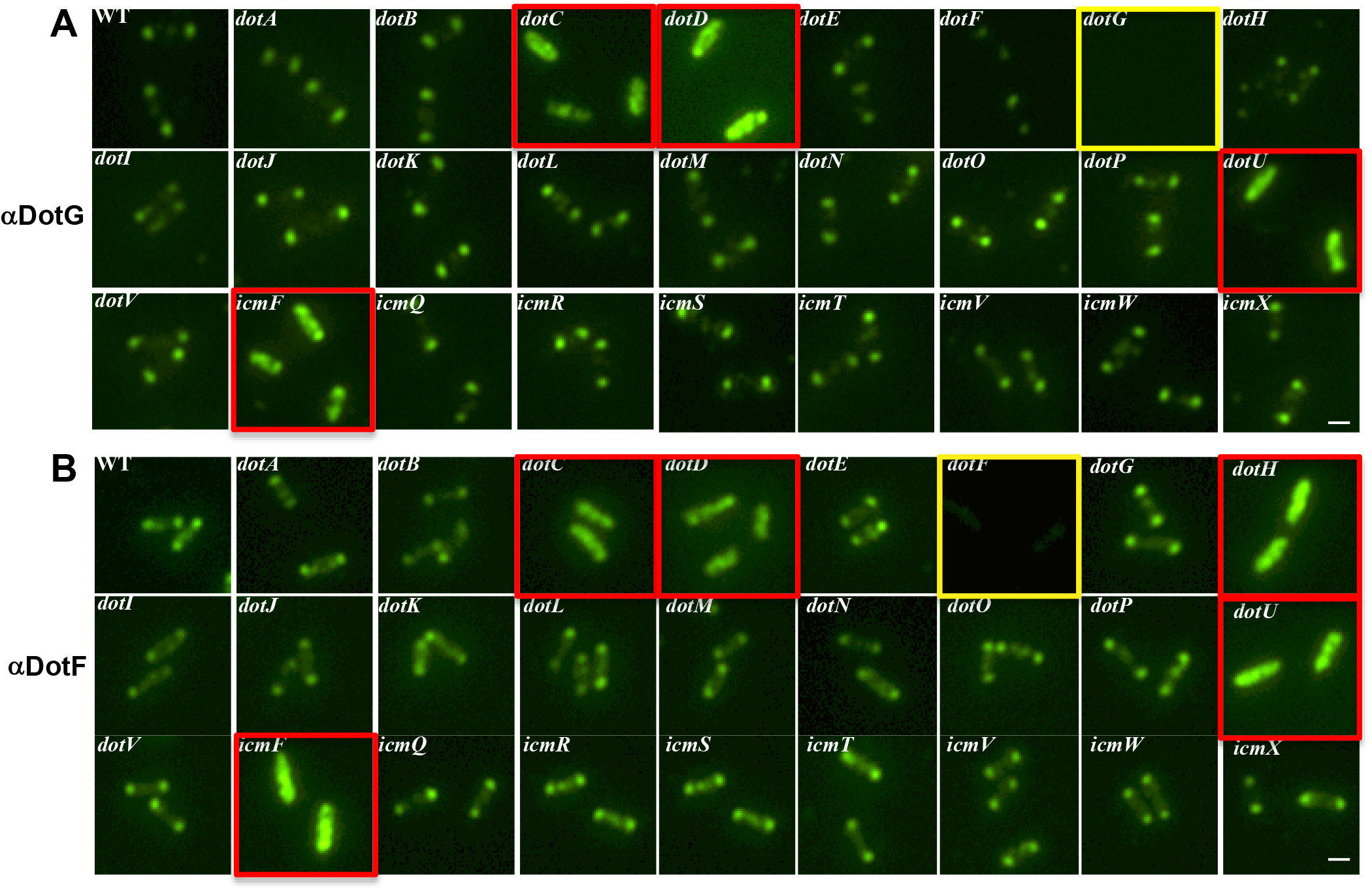
DotG and DotF localization in wild-type *Legionella* and in individual *dot/icm* deletions. DotG and DotF were detected by immunofluorescence microscopy in wild-type cells (WT) and individual *dot/icm* mutant strains. The corresponding deletion strains is boxed in yellow and *dot/icm* deletions that have an effect are boxed in red. Representative images are shown from at least three independent experiments

**Supplementary Fig. 4CD.**
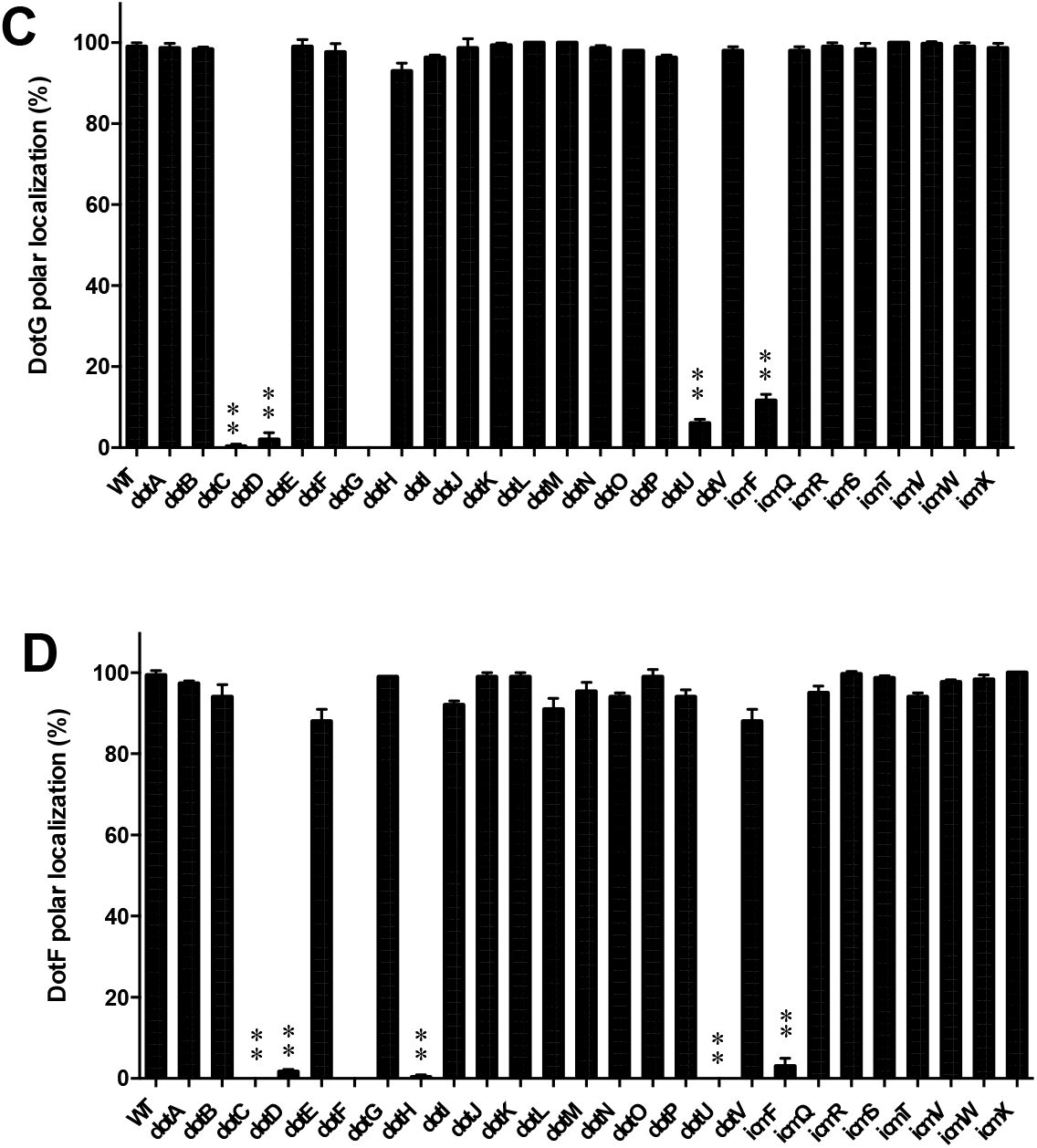
Quantitation of polar localization of DotG and DotF. The percent of cells having polar localization of the Dot proteins in individual *dot/icm* deletions was determined from three independent experiments (at least 100 cells counted from each experiment) and are shown in panels: DotG (C), and DotF (D). Data are presented as means ± SEM. Asterisks indicate statistical difference (** *P*<0.005) compared to the wild-type strain Lp02 (WT) by Student’s *t-*test.

**Supplementary Fig. 5.**
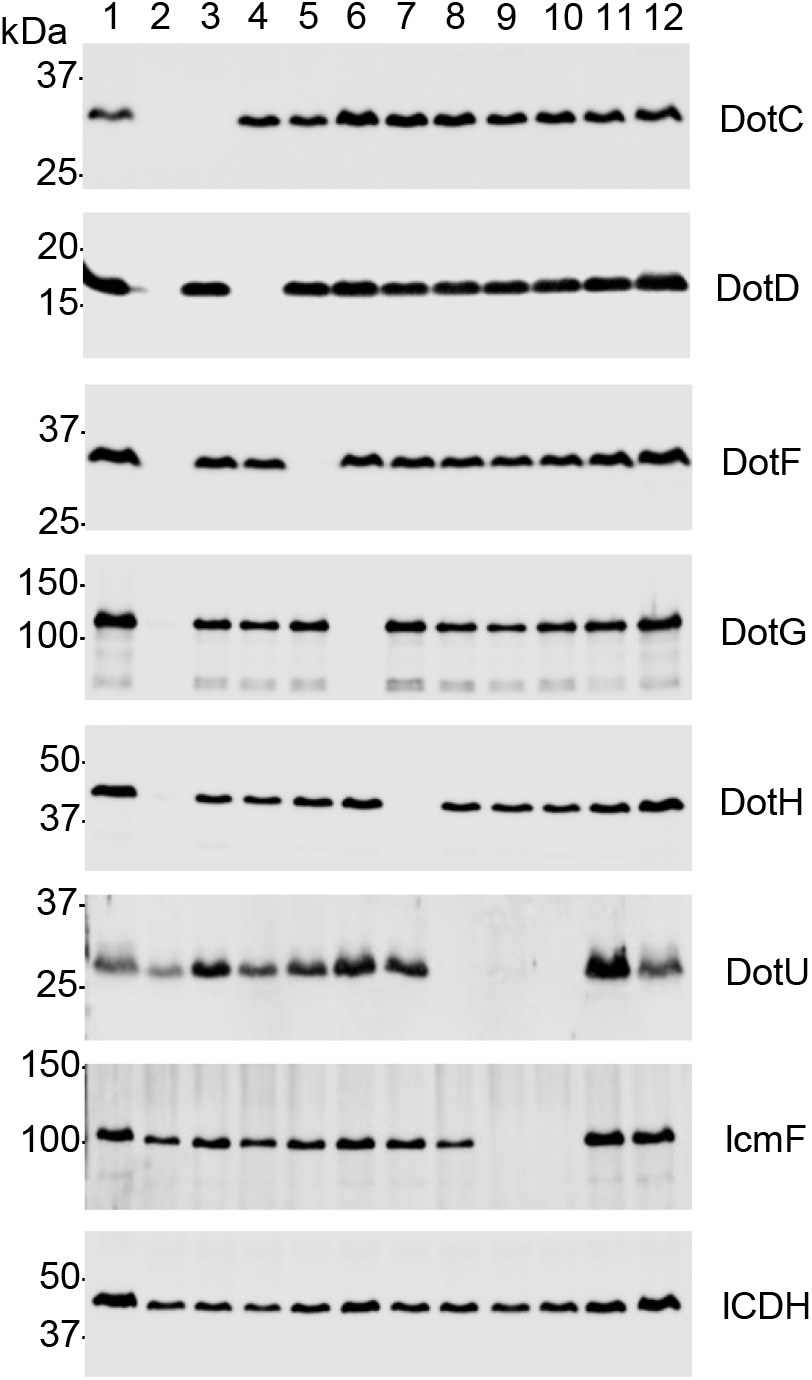
Correct expression of DotC, DotD, DotF, DotG, DotH, DotU and IcmF in various strains in Figure 1B and Supplementary Fig. 4. *L. pneumophila* strains were grown to late exponential phase and westerns were done with the indicated antibodies. ICDH, a cytoplasmic housekeeping protein, was used as a loading control. Samples were loaded in the following order: 1. JV1139 (Lp02 + pJB908), 2. JV5402 (S∆(UF) + vector), 3. JV3743 (∆*dotC*), 4. JV3572 (∆*dotD*), 5. JV3579 (∆*dotF*), 6. JV3559 (∆*dotG*), 7. JV3563 (∆*dotH*), 8. JV4015 (∆*dotU*), 9. JV1179 (∆*icmF*), 10. JV1196 (∆*dotU* ∆*icmF* + vector), 11. JV1199 (∆*dotU* ∆*icmF* + *dotU icmF*), 12. JV1139 (Lp02 + pJB908).

**Supplementary Fig. 6.**
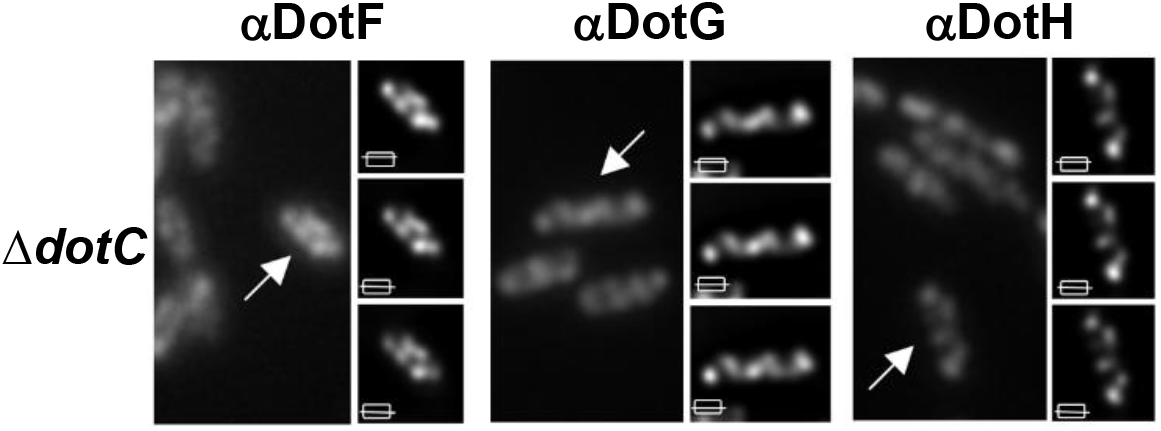
DotH, DotF and DotG proteins are found in non-polar punctae in the ∆*dotC* mutant. DotF, DotG, and DotH localization was assayed using a lower amount of primary antibody and deconvolution microscopy. For each strain, four images are shown including an optical section before deconvolution (left) and three images of a top optical section, a bottom optical section, and a projection of seven Z sections (right panel). Arrows in the left image indicate the cell image that was deconvoluted.

**Supplementary Fig. 7.**
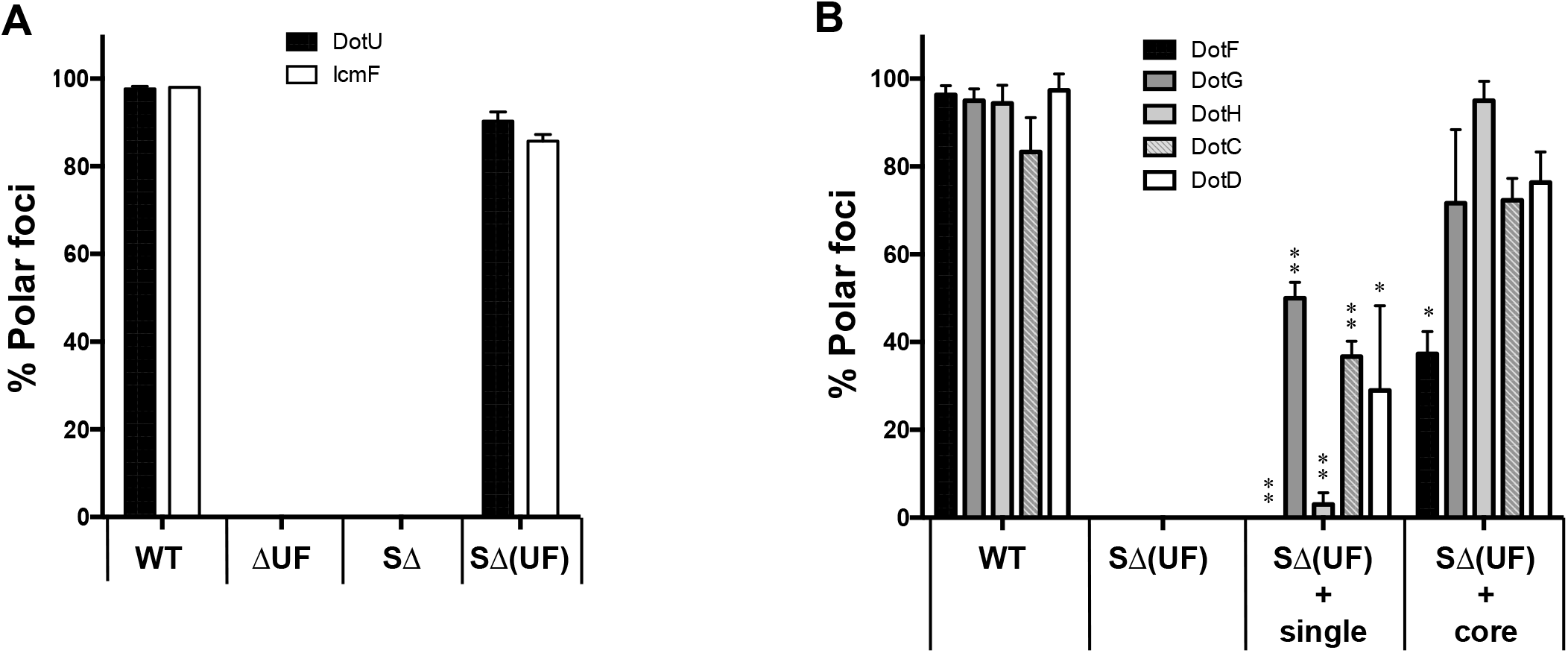
Quantitation of polar localization of Dot proteins for Figure 3. The percent of cells having polar localization of the DotU and IcmF proteins (A) and DotF, DotG, DotH, DotC, and DotD (B) in the wild-type strain Lp02 (WT), ∆*dotU* ∆*icmF* mutant strain (JV1181), the super *dot/icm* deletion strain (S∆, JV4044) and the S∆ strain expressing *dotU* and *icmF* from the chromosome (SΔ(UF), JV5319) was determined from three independent experiments (at least 100 cells counted from each experiment). Data are presented as means ± SEM. Asterisks indicate statistical difference (* *P*<0.05, ** *P*<0.005) compared to the wild-type strain Lp02 (WT) by Student’s *t-*test.

**Supplementary Fig. 8.**
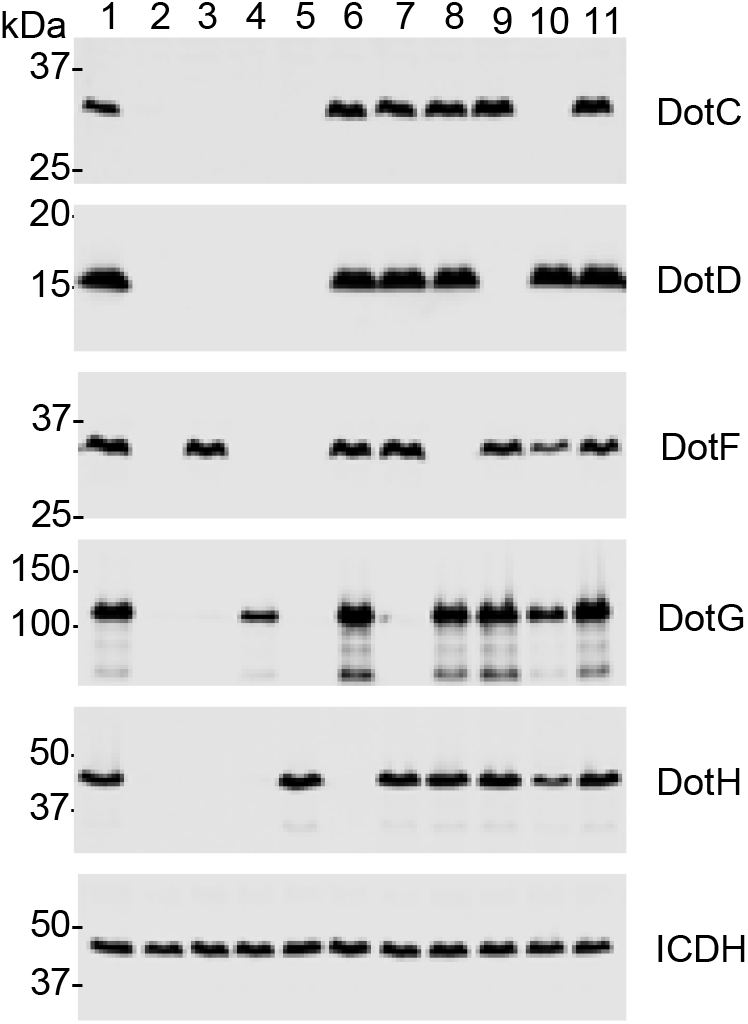
Expression of the correct components in the S∆(UF) strain for Figure 3. *L. pneumophila* strains were grown to late exponential phase and westerns were done with the indicated antibodies. ICDH, a cytoplasmic housekeeping protein, was used as a loading control. Samples were loaded in the following order: 1. JV1139 (Lp02 + pJB908), 2. JV5402 (S∆(UF) + vector), 3. JV5410 (S∆(UF) + *dotC:HA3X*), 4. JV5411 (S∆(UF) + *dotD:HA3X*), 5. JV5403 (S∆(UF) + *dotF*), 6. JV5404 (S∆(UF) + *dotG*), 7. JV5405 (S∆(UF) + *dotH*), 8. JV5443 (S∆(UF) + *dotCDFGH*), 9. JV5750 (S∆(UF) + *dotC:HA3x dotD dotH*), 10. JV5751 (S∆(UF) + *dotD:HA3x dotC dotH*), 11. JV1139 (Lp02 + pJB908).

**Supplementary Fig. 9A.**
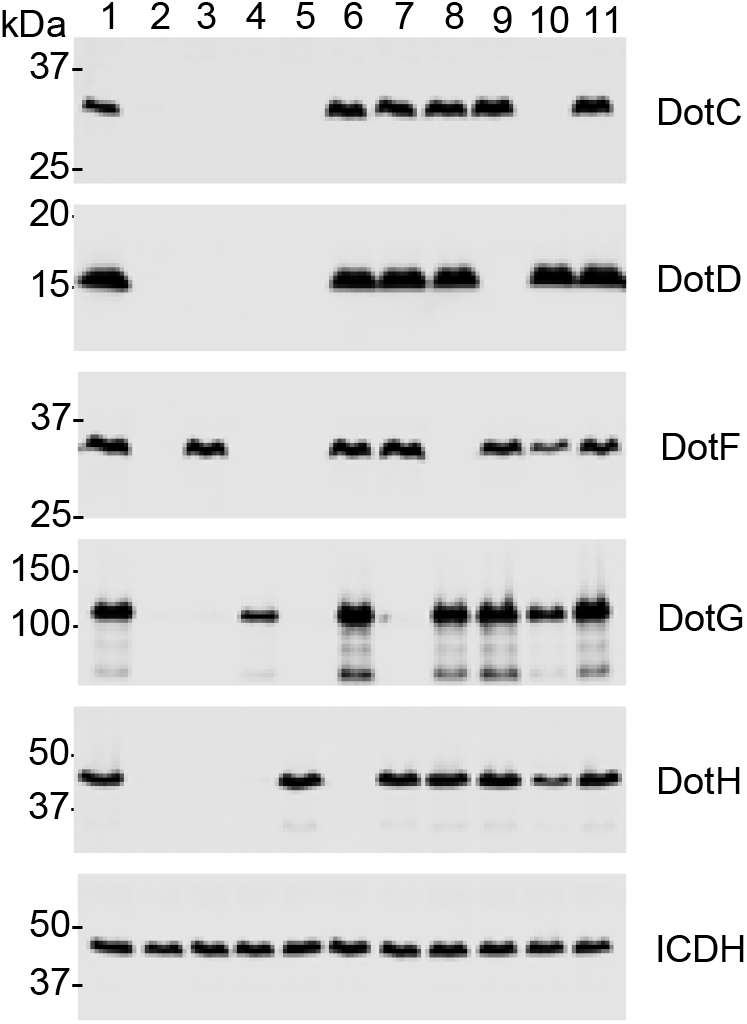
Expression of the correct single and quadruple combinations in the reconstituted S∆(UF) strain for Figure 4. *L. pneumophila* strains were grown to late exponential phase and westerns were done with the indicated antibodies. ICDH, a cytoplasmic housekeeping protein, was used as a loading control. Samples were loaded in the following order: 1. JV1139 (Lp02 + pJB908), 2. JV5402 (S∆(UF) + vector), 3. JV5403 (S∆(UF) + *dotF*), 4. JV5404 (S∆(UF) + *dotG*), 5. JV5405 (S∆(UF) + *dotH*), 6. JV5468 (S∆(UF) + *dotCDFG*), 7. JV5466 (S∆(UF) + *dotCDFH*), 8. JV5475 (S∆(UF) + *dotCDGH*), 9. JV5439 (S∆(UF) + *dotCFGH*), 10. JV5441 (S∆(UF) + *dotDFGH*), 11. JV5443 (S∆(UF) + *dotCDFGH*).

**Supplementary Fig. 9B.**
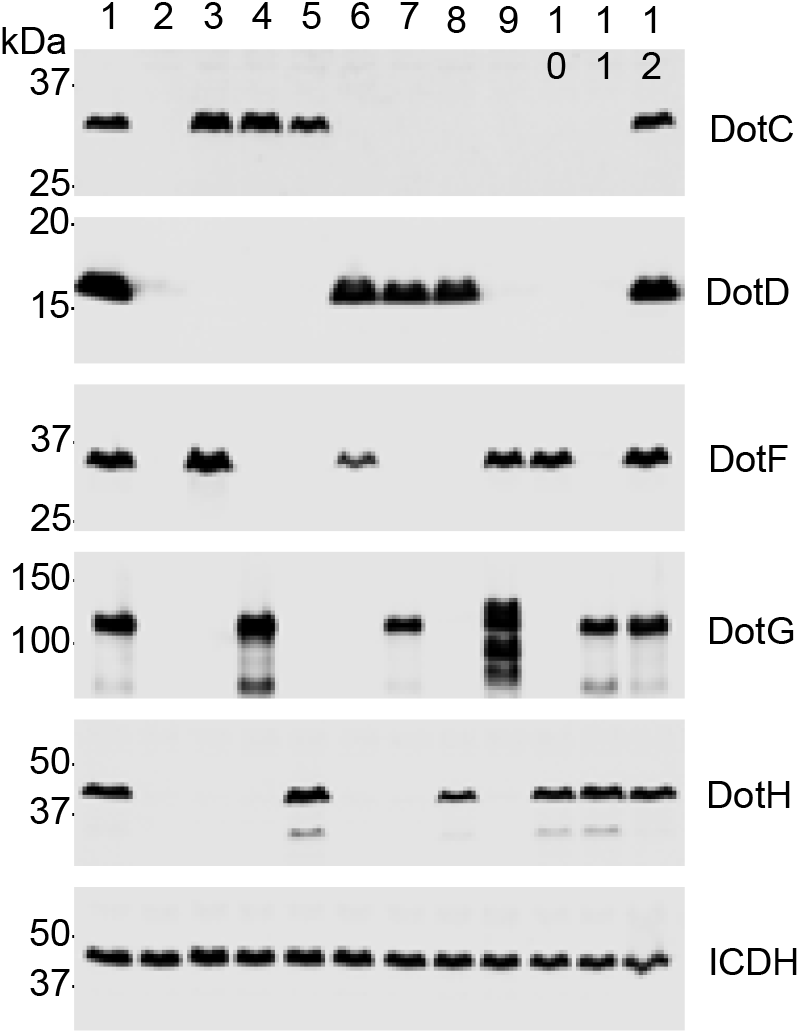
Expression of the correct double combinations in the reconstituted S∆(UF) strain for Figure 4. *L. pneumophila* strains were grown to late exponential phase and westerns were done with the indicated antibodies. ICDH, a cytoplasmic housekeeping protein, was used as a loading control. Samples were loaded in the following order: 1. JV1139 (Lp02 + pJB908), 2. JV5402 (S∆(UF) + vector), 3. JV5452 (S∆(UF) + *dotCF*), 4. JV5455 (S∆(UF) + *dotCG*), 5. JV5458 (S∆(UF) + *dotCH*), 6. JV5453 (S∆(UF) + *dotDF*), 7. JV5456 (S∆(UF) + *dotDG*), 8. JV5459 (S∆(UF) + *dotDH*), 9. JV5408 (S∆(UF) + *dotFG*), 10. JV5407 (S∆(UF) + *dotFH*), 11. JV5406 (S∆(UF) + *dotGH*), 12. JV1139 (Lp02 + pJB908).

**Supplementary Fig. 9C.**
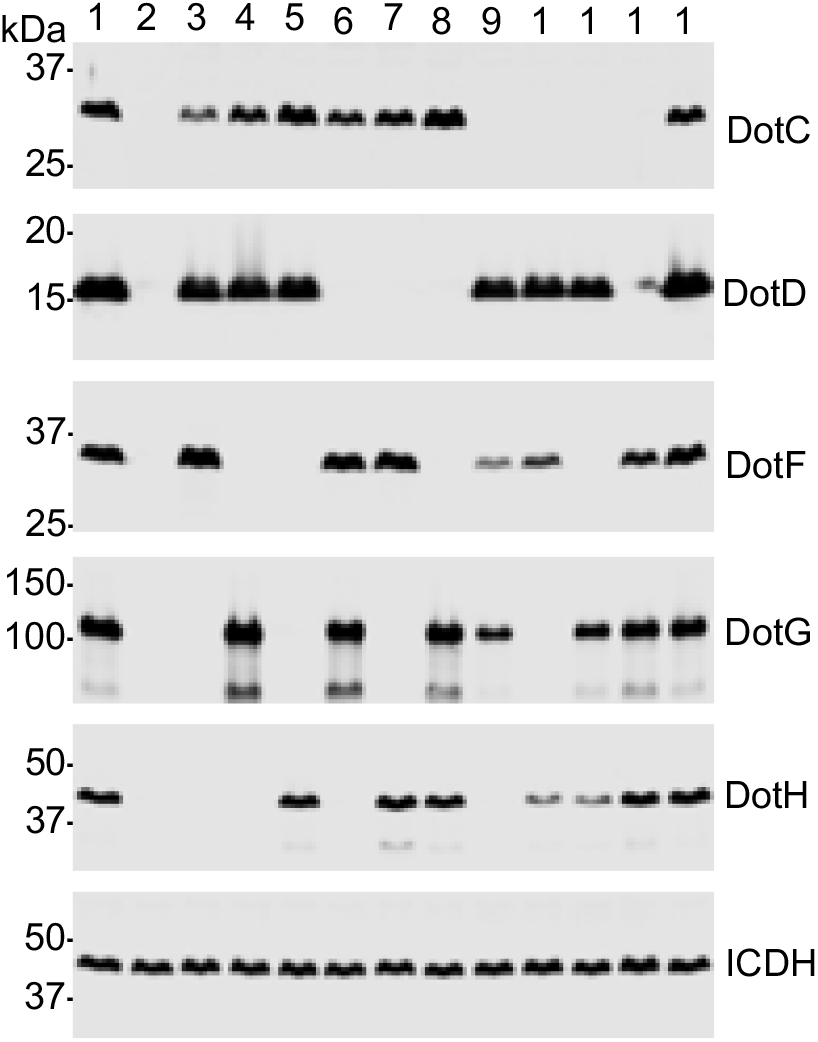
Expression of the correct triple combinations in the reconstituted S∆(UF) strain for Figure 4. *L. pneumophila* strains were grown to late exponential phase and westerns were done with the indicated antibodies. ICDH, a cytoplasmic housekeeping protein, was used as a loading control. Samples were loaded in the following order: 1. JV1139 (Lp02 + pJB908), 2. JV5402 (S∆(UF) + vector), 3. JV5454 (S∆(UF) + *dotCDF*), 4. JV5457 (S∆(UF) + *dotCDG*), 5. JV5460 (S∆(UF) + *dotCDH*), 6. JV5467 (S∆(UF) + *dotCFG*), 7. JV5464 (S∆(UF) + *dotCFH*), 8. JV5473 (S∆(UF) + *dotCGH*), 9. JV5472 (S∆(UF) + *dotDFG*), 10. JV5465 (S∆(UF) + *dotDFH*), 11. JV5474 (S∆(UF) + *dotDGH*), 12. JV5409 (S∆(UF) + *dotFGH*), 13. JV1139 (Lp02 + pJB908).

**Supplementary Fig. 9D.**
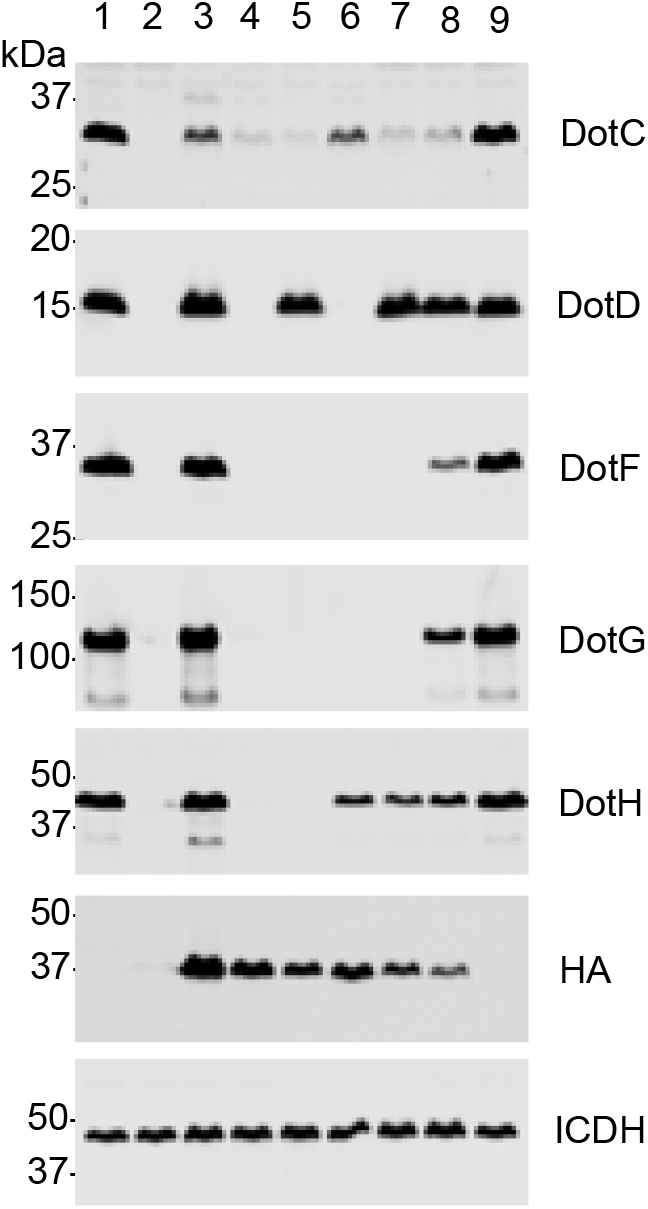
Expression of the correct components in the reconstituted S∆(UF) strain containing HA-tagged proteins for Figure 4. *L. pneumophila* strains were grown to late exponential phase and westerns were done with the indicated antibodies. ICDH, a cytoplasmic housekeeping protein, was used as a loading control. Samples were loaded in the following order: 1. JV1139 (Lp02 + pJB908), 2. JV5402 (S∆(UF) + vector), 3. JV5484 (∆*dotC* + *dotC:HA3x*), 4. JV5480 (S∆(UF) + *dotC:HA3x*), 5. JV5482 (S∆(UF) + *dotC:HA3x dotD*), 6. JV5749 (S∆(UF) + *dotC:HA3x dotH*), 7. JV5750 (S∆(UF) + *dotC:HA3x dotD dotH*), 8. JV5752 (S∆(UF) + *dotC:HA3x dotD dotFGH*), 9. JV1139 (Lp02 + pJB908).

**Supplementary Fig. 9E.**
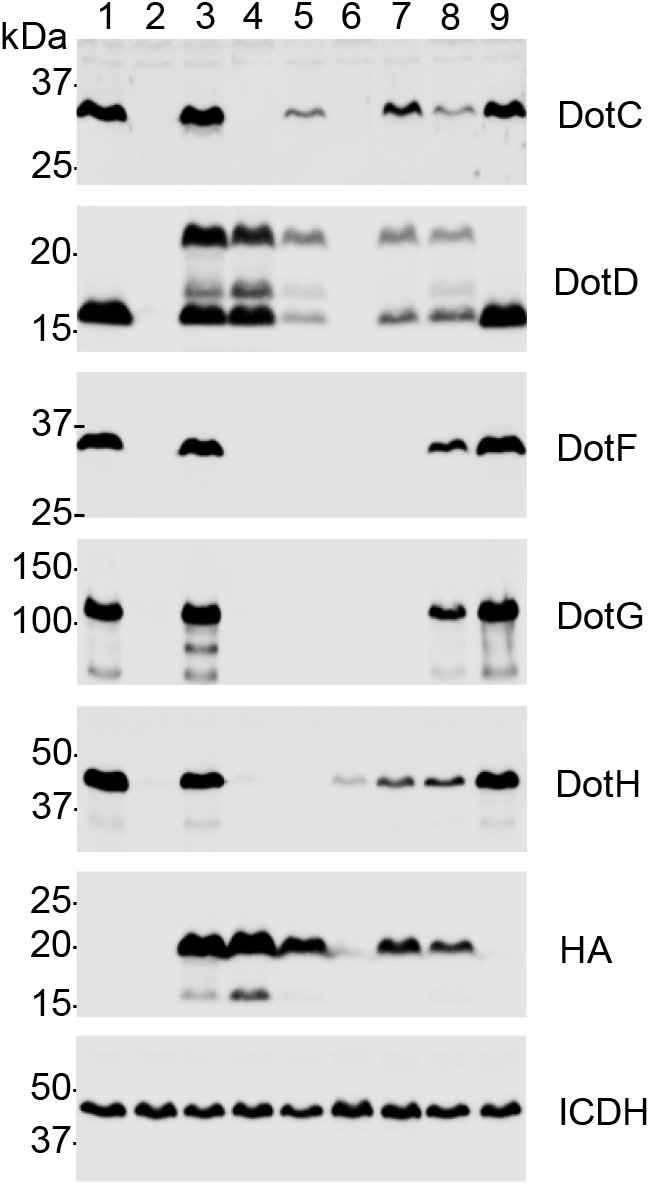
Expression of the correct components in the reconstituted S∆(UF) strain containing HA-tagged proteins for Figure 4. *L. pneumophila* strains were grown to late exponential phase and westerns were done with the indicated antibodies. ICDH, a cytoplasmic housekeeping protein, was used as a loading control. Samples were loaded in the following order: 1. JV1139 (Lp02 + pJB908), 2. JV5402 (S∆(UF) + vector), 3. JV5484 (∆*dotD* + *dotD:HA3x*), 4. JV5480 (S∆(UF) + *dotD:HA3x*), 5. JV5482 (S∆(UF) + *dotD:HA3x dotC*), 6. JV5749 (S∆(UF) + *dotD:HA3x dotH*), 7. JV5750 (S∆(UF) + *dotD:HA3x dotC dotH*), 8. JV5752 (S∆(UF) + *dotD:HA3x dotC dotFGH*), 9. JV1139 (Lp02 + pJB908).

**Supplementary Fig. 10.**
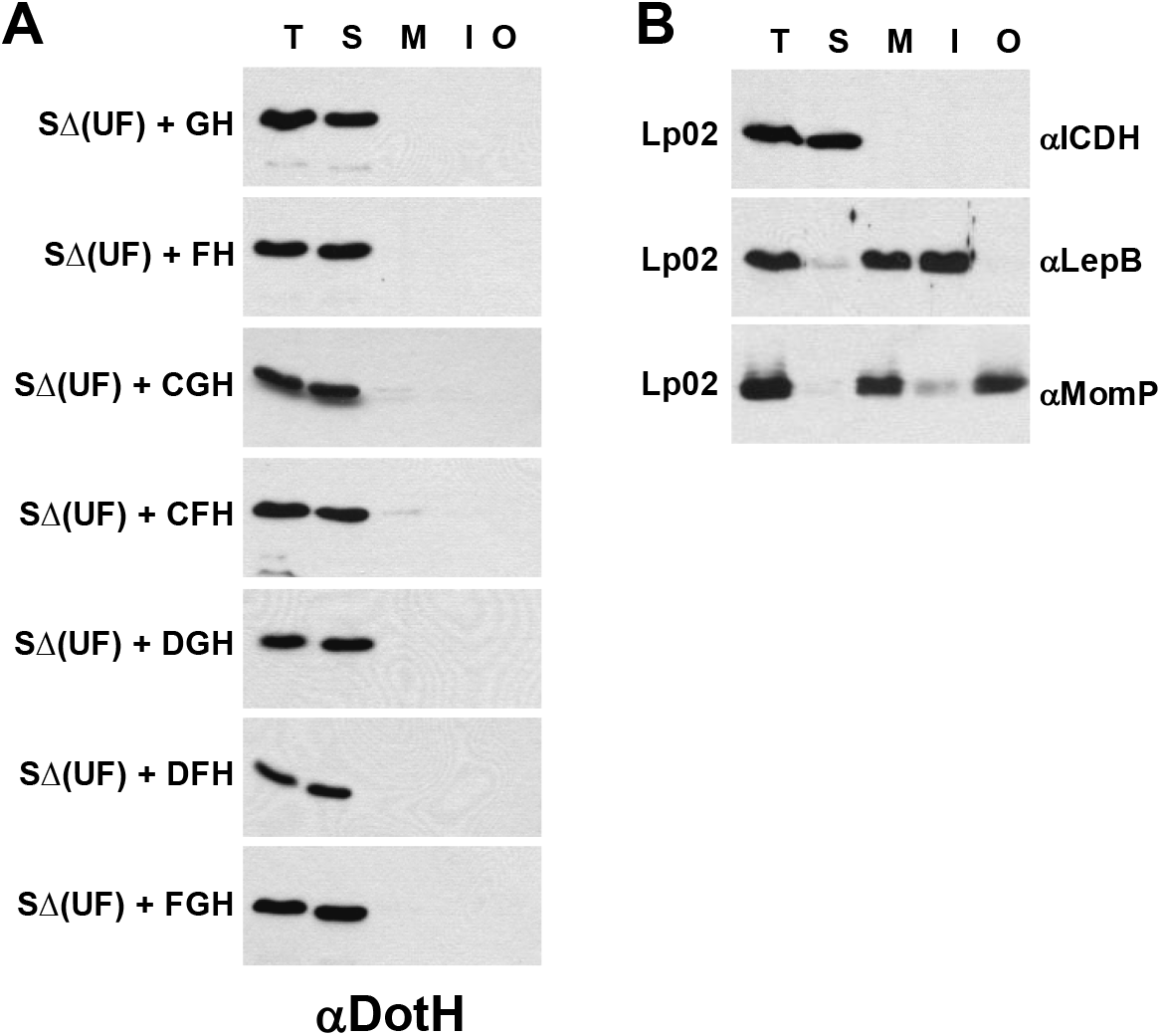
DotH localization to the outer membrane requires the anchor proteins DotU and IcmF. Cells were fractionated by a combination of ultracentrifugation and Triton X-100 solubility, proteins were separated by SDS-PAGE and probed in Westerns using DotH specific antibodies. (A) S∆(UF) strains that did not restore DotH outer membrane localization include: DotG/DotH (JV5406), DotF/DotH (JV5407), DotC/DotG/DotH (JV5460), DotC/DotF/DotH (JV5464), DotG/DotF/DotH (JV5409), DotD/DotF/DotH (JV5465) and DotF/DotG/DotH (JV5409). (B). Controls for the fractionations are shown and include the cytoplasmic protein isocitrate dehydrogenase (ICDH), the inner membrane protein LepB and the outer membrane protein MomP. Experiments were done in triplicate and representative images are shown.

**Supplementary Fig. 11.**
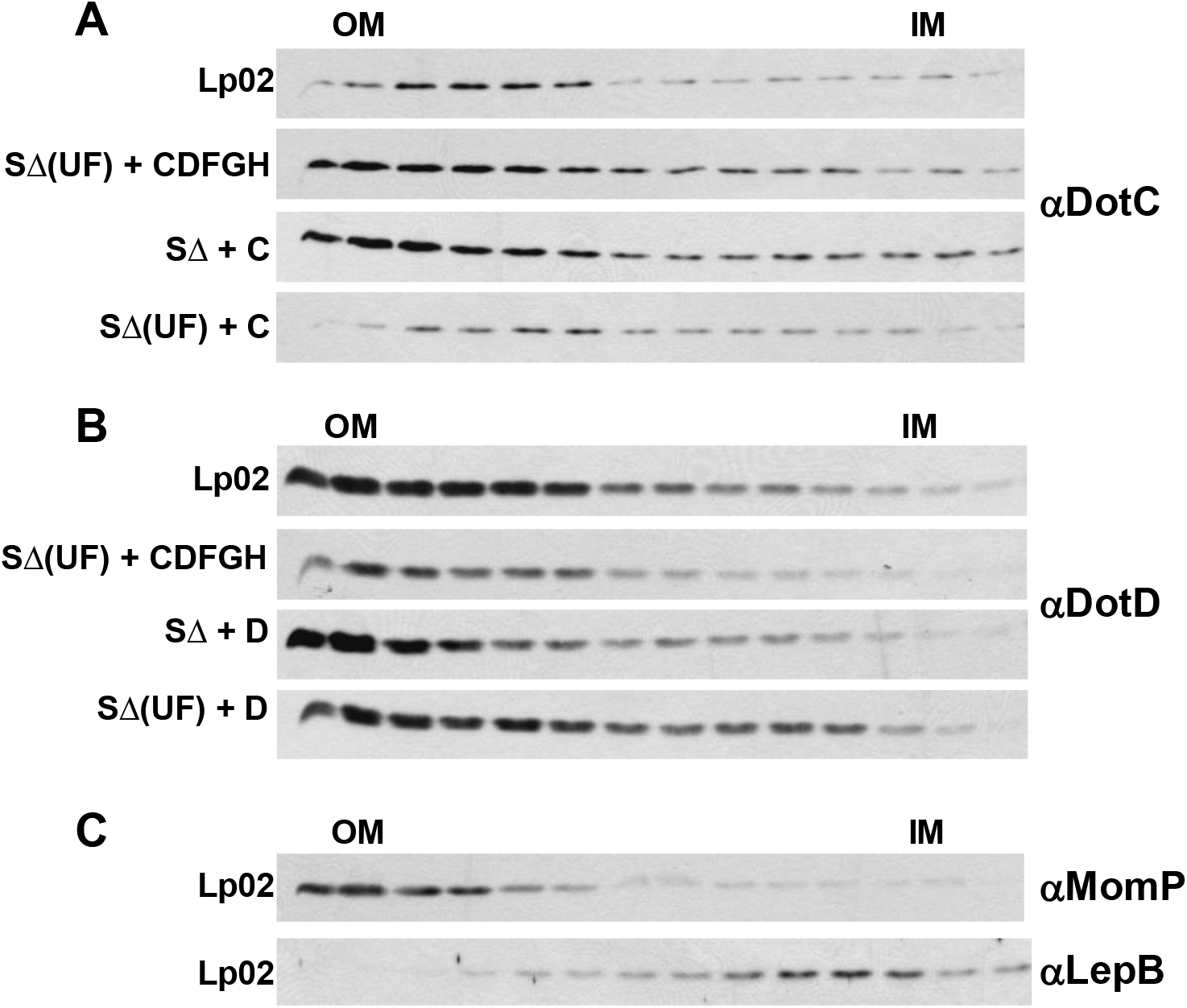
Lipoproteins DotC and DotD remain in the outer membrane in the presence and absence of DotU and IcmF. Strains were grown to late exponential phase, total membrane was isolated and separated by sucrose density gradient. Fractions were then separated by SDS-PAGE and probed using DotC or DotD specific antibodies (indicated to the right of the blots). (A) Wild-type *Legionella* Lp02, S∆(UF) + core (JV5443), S∆ + DotC (JV4225), and S∆(UF) + DotC (JV5410). (B). Wild-type *Legionella* Lp02, S∆(UF) + core (JV5443), S∆ + DotD (JV4695), and S∆(UF) + DotD (JV5411). (C). Controls include the outer membrane protein MomP and the inner membrane protein LepB.

**Supplementary Table 1.**
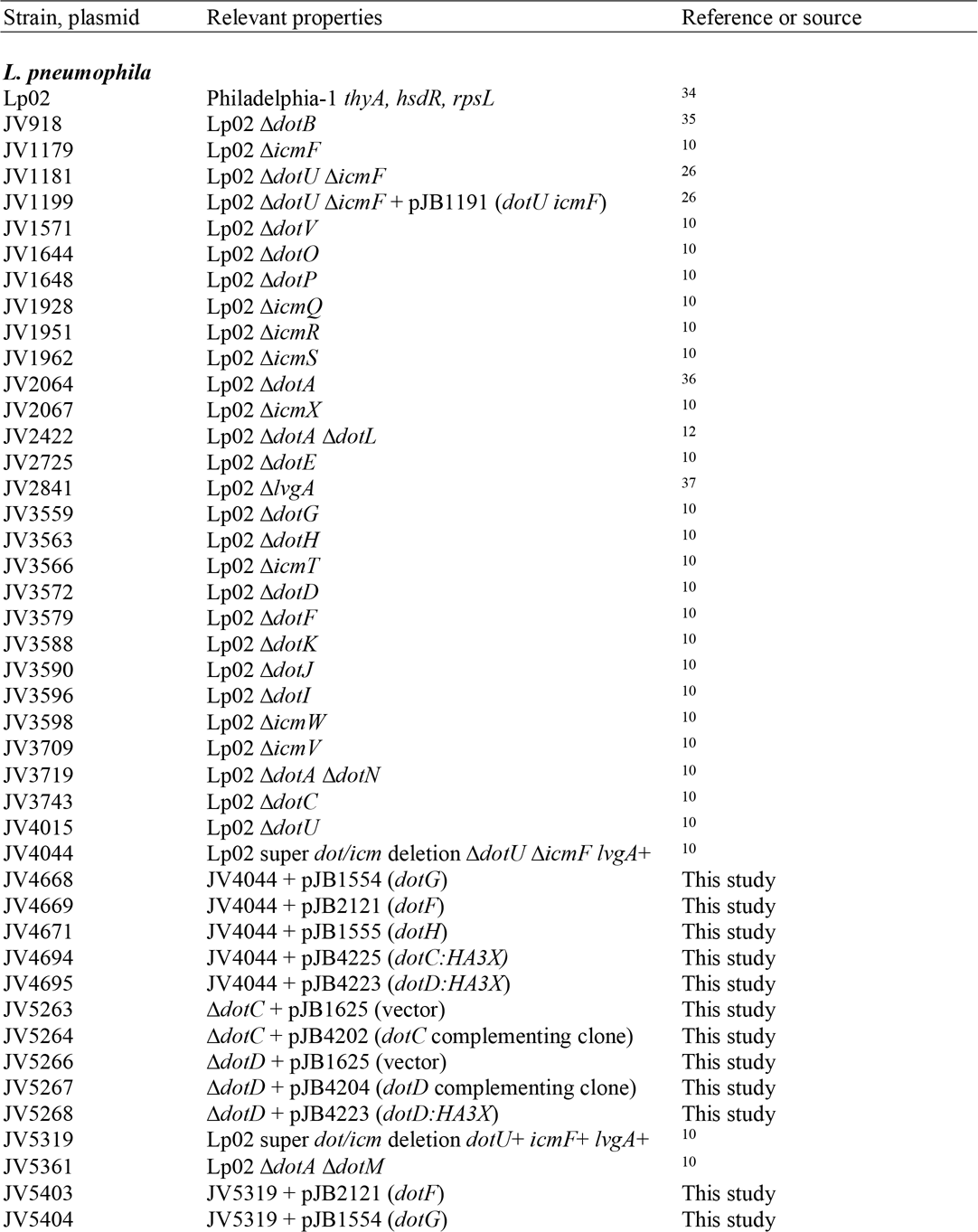
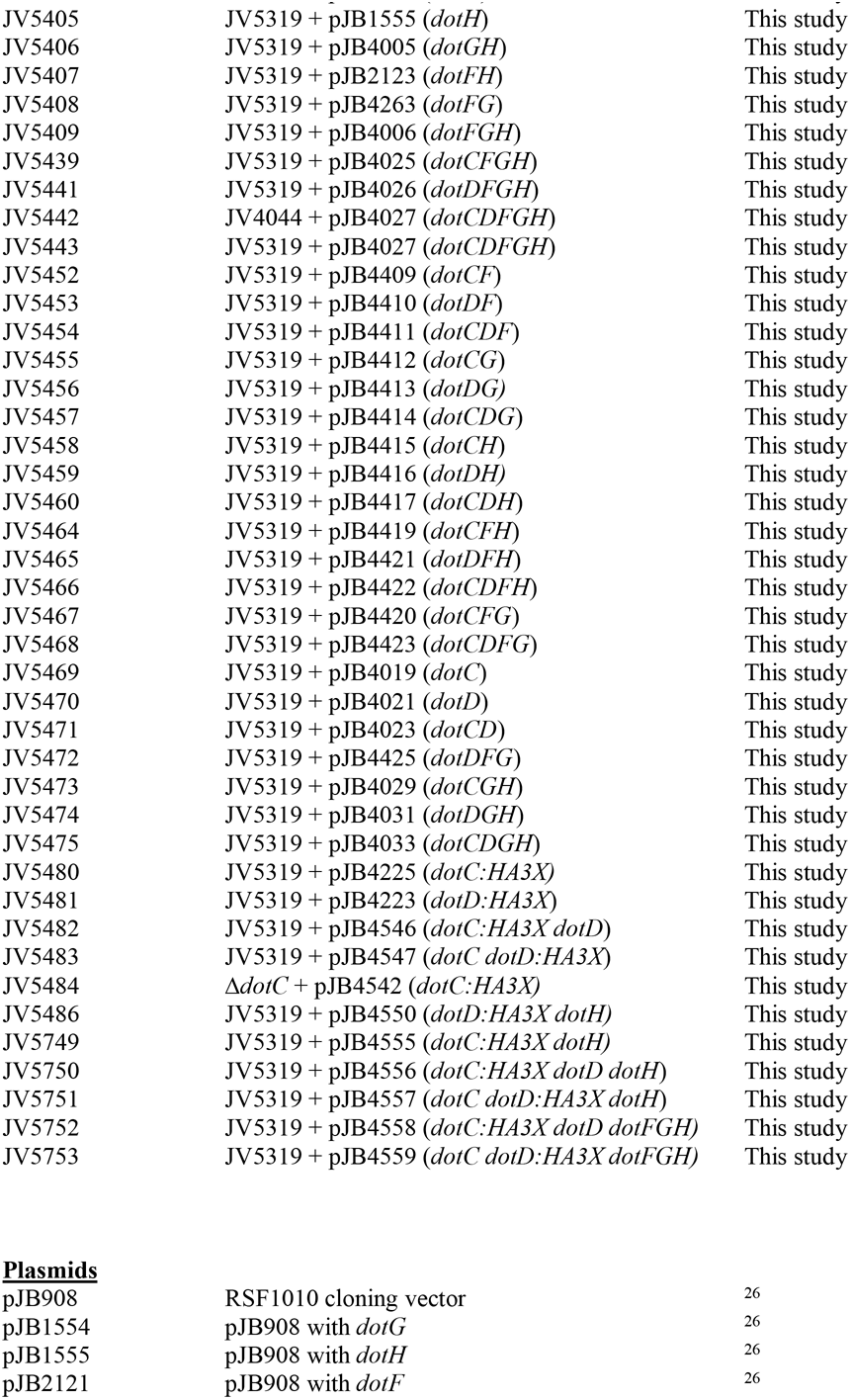
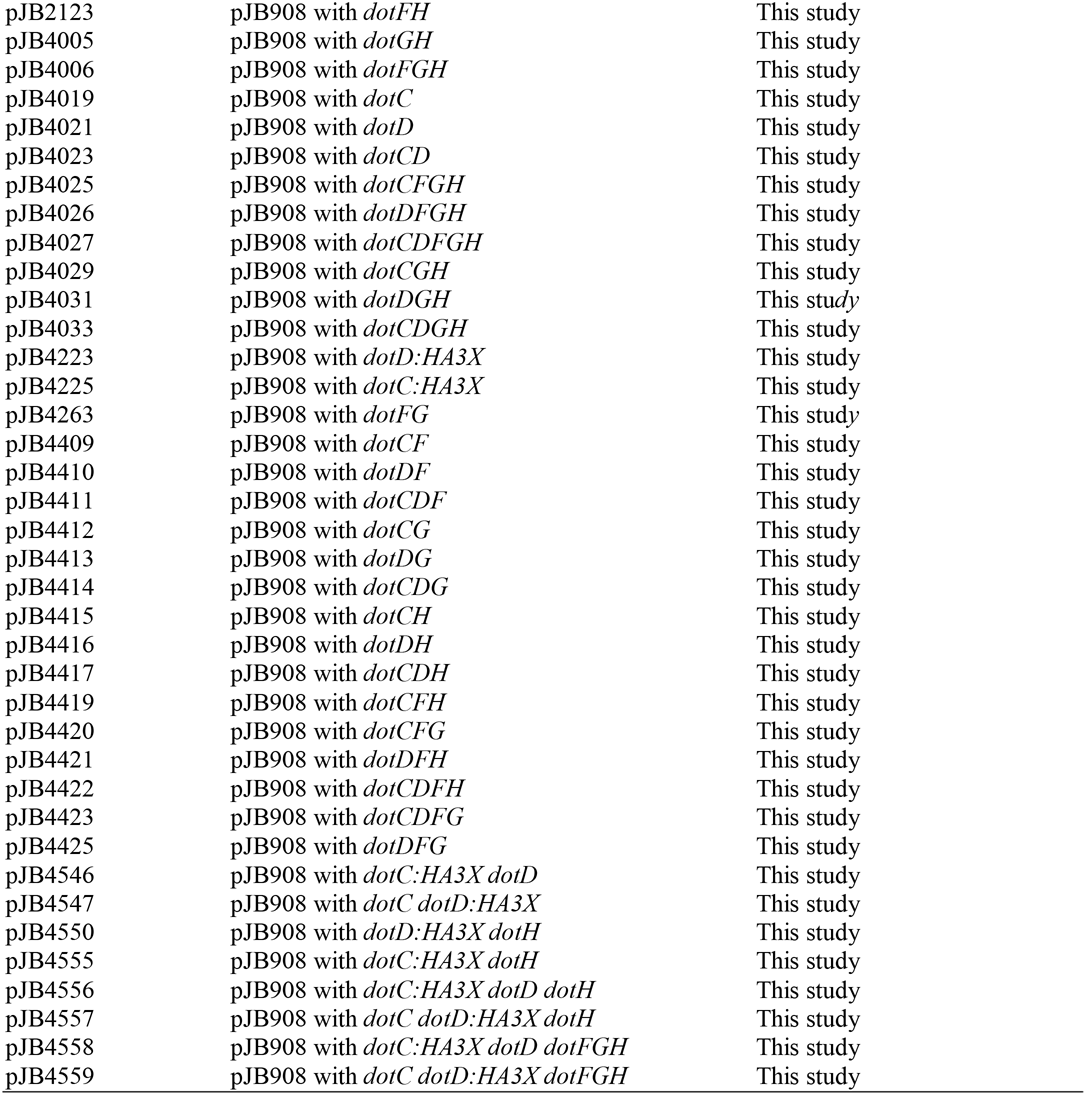
Bacterial strains, plasmids, and primers employed in this study

| Strain, plasmid | Relevant properties | Reference or source |
| --- | --- | --- |
| ***L. pneumophila*** |  |  |
| Lp02 | Philadelphia-1 *thyA*, *hsdR*, *rpsL* | 34 |
| JV918 | Lp02 $\Delta dotB$ | 35 |
| JV1179 | Lp02 $\Delta icmF$ | 10 |
| JV1181 | Lp02 $\Delta dotU \Delta icmF$ | 26 |
| JV1199 | Lp02 $\Delta dotU \Delta icmF$ + pJB1191 (*dotU icmF*) | 26 |
| JV1571 | Lp02 $\Delta dotV$ | 10 |
| JV1644 | Lp02 $\Delta dotO$ | 10 |
| JV1648 | Lp02 $\Delta dotP$ | 10 |
| JV1928 | Lp02 $\Delta icmQ$ | 10 |
| JV1951 | Lp02 $\Delta icmR$ | 10 |
| JV1962 | Lp02 $\Delta icmS$ | 10 |
| JV2064 | Lp02 $\Delta dotA$ | 36 |
| JV2067 | Lp02 $\Delta icmX$ | 10 |
| JV2422 | Lp02 $\Delta dotA \Delta dotL$ | 12 |
| JV2725 | Lp02 $\Delta dotE$ | 10 |
| JV2841 | Lp02 $\Delta lvgA$ | 37 |
| JV3559 | Lp02 $\Delta dotG$ | 10 |
| JV3563 | Lp02 $\Delta dotH$ | 10 |
| JV3566 | Lp02 $\Delta icmT$ | 10 |
| JV3572 | Lp02 $\Delta dotD$ | 10 |
| JV3579 | Lp02 $\Delta dotF$ | 10 |
| JV3588 | Lp02 $\Delta dotK$ | 10 |
| JV3590 | Lp02 $\Delta dotJ$ | 10 |
| JV3596 | Lp02 $\Delta dotI$ | 10 |
| JV3598 | Lp02 $\Delta icmW$ | 10 |
| JV3709 | Lp02 $\Delta icmV$ | 10 |
| JV3719 | Lp02 $\Delta dotA \Delta dotN$ | 10 |
| JV3743 | Lp02 $\Delta dotC$ | 10 |
| JV4015 | Lp02 $\Delta dotU$ | 10 |
| JV4044 | Lp02 super *dot/icm* deletion $\Delta dotU \Delta icmF lvgA+$ | 10 |
| JV4668 | JV4044 + pJB1554 (*dotG*) | This study |
| JV4669 | JV4044 + pJB2121 (*dotF*) | This study |
| JV4671 | JV4044 + pJB1555 (*dotH*) | This study |
| JV4694 | JV4044 + pJB4225 (*dotC:HA3X*) | This study |
| JV4695 | JV4044 + pJB4223 (*dotD:HA3X*) | This study |
| JV5263 | $\Delta dotC$ + pJB1625 (vector) | This study |
| JV5264 | $\Delta dotC$ + pJB4202 (*dotC* complementing clone) | This study |
| JV5266 | $\Delta dotD$ + pJB1625 (vector) | This study |
| JV5267 | $\Delta dotD$ + pJB4204 (*dotD* complementing clone) | This study |
| JV5268 | $\Delta dotD$ + pJB4223 (*dotD:HA3X*) | This study |
| JV5319 | Lp02 super *dot/icm* deletion *dotU+ icmF+ lvgA+* | 10 |
| JV5361 | Lp02 $\Delta dotA \Delta dotM$ | 10 |
| JV5403 | JV5319 + pJB2121 (*dotF*) | This study |
| JV5404 | JV5319 + pJB1554 (*dotG*) | This study |
| JV5405 | JV5319 + pJB1555 (*dotH*) | This study |
| JV5406 | JV5319 + pJB4005 (*dotGH*) | This study |
| JV5407 | JV5319 + pJB2123 (*dotFH*) | This study |
| JV5408 | JV5319 + pJB4263 (*dotFG*) | This study |
| JV5409 | JV5319 + pJB4006 (*dotFGH*) | This study |
| JV5439 | JV5319 + pJB4025 (*dotCFGH*) | This study |
| JV5441 | JV5319 + pJB4026 (*dotDFGH*) | This study |
| JV5442 | JV4044 + pJB4027 (*dotCDFGH*) | This study |
| JV5443 | JV5319 + pJB4027 (*dotCDFGH*) | This study |
| JV5452 | JV5319 + pJB4409 (*dotCF*) | This study |
| JV5453 | JV5319 + pJB4410 (*dotDF*) | This study |
| JV5454 | JV5319 + pJB4411 (*dotCDF*) | This study |
| JV5455 | JV5319 + pJB4412 (*dotCG*) | This study |
| JV5456 | JV5319 + pJB4413 (*dotDG*) | This study |
| JV5457 | JV5319 + pJB4414 (*dotCDG*) | This study |
| JV5458 | JV5319 + pJB4415 (*dotCH*) | This study |
| JV5459 | JV5319 + pJB4416 (*dotDH*) | This study |
| JV5460 | JV5319 + pJB4417 (*dotCDH*) | This study |
| JV5464 | JV5319 + pJB4419 (*dotCFH*) | This study |
| JV5465 | JV5319 + pJB4421 (*dotDFH*) | This study |
| JV5466 | JV5319 + pJB4422 (*dotCDFH*) | This study |
| JV5467 | JV5319 + pJB4420 (*dotCFG*) | This study |
| JV5468 | JV5319 + pJB4423 (*dotCDFG*) | This study |
| JV5469 | JV5319 + pJB4019 (*dotC*) | This study |
| JV5470 | JV5319 + pJB4021 (*dotD*) | This study |
| JV5471 | JV5319 + pJB4023 (*dotCD*) | This study |
| JV5472 | JV5319 + pJB4425 (*dotDFG*) | This study |
| JV5473 | JV5319 + pJB4029 (*dotCGH*) | This study |
| JV5474 | JV5319 + pJB4031 (*dotDGH*) | This study |
| JV5475 | JV5319 + pJB4033 (*dotCDGH*) | This study |
| JV5480 | JV5319 + pJB4225 (*dotC:HA3X*) | This study |
| JV5481 | JV5319 + pJB4223 (*dotD:HA3X*) | This study |
| JV5482 | JV5319 + pJB4546 (*dotC:HA3X dotD*) | This study |
| JV5483 | JV5319 + pJB4547 (*dotC dotD:HA3X*) | This study |
| JV5484 | $\Delta dotC$ + pJB4542 (*dotC:HA3X*) | This study |
| JV5486 | JV5319 + pJB4550 (*dotD:HA3X dotH*) | This study |
| JV5749 | JV5319 + pJB4555 (*dotC:HA3X dotH*) | This study |
| JV5750 | JV5319 + pJB4556 (*dotC:HA3X dotD dotH*) | This study |
| JV5751 | JV5319 + pJB4557 (*dotC dotD:HA3X dotH*) | This study |
| JV5752 | JV5319 + pJB4558 (*dotC:HA3X dotD dotFGH*) | This study |
| JV5753 | JV5319 + pJB4559 (*dotC dotD:HA3X dotFGH*) | This study |
| **Plasmids** |  |  |
| pJB908 | RSF1010 cloning vector | 26 |
| pJB1554 | pJB908 with *dotG* | 26 |
| pJB1555 | pJB908 with *dotH* | 26 |
| pJB2121 | pJB908 with *dotF* | 26 |
| pJB2123 | pJB908 with *dotFH* | This study |
| pJB4005 | pJB908 with *dotGH* | This study |
| pJB4006 | pJB908 with *dotFGH* | This study |
| pJB4019 | pJB908 with *dotC* | This study |
| pJB4021 | pJB908 with *dotD* | This study |
| pJB4023 | pJB908 with *dotCD* | This study |
| pJB4025 | pJB908 with *dotCFGH* | This study |
| pJB4026 | pJB908 with *dotDFGH* | This study |
| pJB4027 | pJB908 with *dotCDFGH* | This study |
| pJB4029 | pJB908 with *dotCGH* | This study |
| pJB4031 | pJB908 with *dotDGH* | This study |
| pJB4033 | pJB908 with *dotCDGH* | This study |
| pJB4223 | pJB908 with *dotD:HA3X* | This study |
| pJB4225 | pJB908 with *dotC:HA3X* | This study |
| pJB4263 | pJB908 with *dotFG* | This study |
| pJB4409 | pJB908 with *dotCF* | This study |
| pJB4410 | pJB908 with *dotDF* | This study |
| pJB4411 | pJB908 with *dotCDF* | This study |
| pJB4412 | pJB908 with *dotCG* | This study |
| pJB4413 | pJB908 with *dotDG* | This study |
| pJB4414 | pJB908 with *dotCDG* | This study |
| pJB4415 | pJB908 with *dotCH* | This study |
| pJB4416 | pJB908 with *dotDH* | This study |
| pJB4417 | pJB908 with *dotCDH* | This study |
| pJB4419 | pJB908 with *dotCFH* | This study |
| pJB4420 | pJB908 with *dotCFG* | This study |
| pJB4421 | pJB908 with *dotDFH* | This study |
| pJB4422 | pJB908 with *dotCDFH* | This study |
| pJB4423 | pJB908 with *dotCDFG* | This study |
| pJB4425 | pJB908 with *dotDFG* | This study |
| pJB4546 | pJB908 with *dotC:HA3X dotD* | This study |
| pJB4547 | pJB908 with *dotC dotD:HA3X* | This study |
| pJB4550 | pJB908 with *dotD:HA3X dotH* | This study |
| pJB4555 | pJB908 with *dotC:HA3X dotH* | This study |
| pJB4556 | pJB908 with *dotC:HA3X dotD dotH* | This study |
| pJB4557 | pJB908 with *dotC dotD:HA3X dotH* | This study |
| pJB4558 | pJB908 with *dotC:HA3X dotD dotFGH* | This study |
| pJB4559 | pJB908 with *dotC dotD:HA3X dotFGH* | This study |

